# Compensatory mechanisms render Tcf7l1a dispensable for eye formation despite its cell-autonomous requirement in eye field specification

**DOI:** 10.1101/377119

**Authors:** Rodrigo M. Young, Florencia Cavodeassi, Thomas A. Hawkins, Heather L. Stickney, Quenten Schwarz, Lisa M. Lawrence, Claudia Wierzbicki, Gaia Gestri, Elizabeth Ambrosio, Allison Klosner, Jasmine Rowell, Isaac H. Bianco, Miguel L. Allende, Stephen W. Wilson

## Abstract

The vertebrate eye originates from the eyefield, a domain of cells specified by a small number of transcription factors. In this study, we show that Tcf7la is one such transcription factor that acts cell-autonomously to specify the eye field in zebrafish. Despite the much reduced eyefield in *tcf7l1a* mutants, these fish develop normal eyes revealing a striking ability of the eye to recover from a severe early phenotype. This robustness is not mediated through compensation by paralogous genes; instead, the smaller optic vesicle of *tcf7l1a* mutants shows delayed neurogenesis and continues to grow until it achieves approximately normal size. Although the developing eye is robust to the lack of Tcf7l1a function, it is sensitised to the effects of additional mutations. In support of this, a forward genetic screen identified mutations in *hesx1, cct5* and *gdf6a*, which give synthetically enhanced eye specification or growth phenotypes when in combination with the *tcf7l1a* mutation.

## Introduction

The paired optic vesicles originate from the eye field, a single, coherent group of cells located in the anterior neural plate (Cavodeassi 2018). During early neural development, the specification and relative sizes of prospective forebrain territories, including the eye field, depend on the activity of the Wnt/β-Catenin and other signalling pathways (Beccari et al., 2013; Cavodeassi 2014; Wilson and Houart, 2004). Enhanced Wnt/β-Catenin activity leads to embryos with small or no eyes (Cavodeassi et al., 2005; Kim et al., 2000; Heisenberg at al., 2001; Houart et al., 2002). In contrast, decreasing activity of Wnt/β Catenin signalling generates embryos with bigger forebrain and eyes (Cavodeassi et al., 2005; Glinka et al., 1998; Lekven et al., 2001; Houart et al., 2002). Although much research has focused on the molecular mechanisms involved in the specification of the eye field, little is known about what happens to the eyes if eye field size is disrupted.

A number of genes have been identified as encoding a transcription factor network that specifies the eye field (Beccari et al 2013; Zuber et al. 2003). These genes have been defined based on conserved cross species expression patterns in the anterior neuroectoderm and on phenotypes observed when overexpressed or when function is compromised (Beccari et al., 2013). Perhaps surprisingly, to date there are relatively few mutations that lead to complete loss of eyes suggesting that early stages of eye development are robust to compromised function of genes involved in eye development. Indeed, in humans, eye phenotypes are often highly variable in terms of penetrance and expressivity even between left and right eyes (Reis and Semina, 2015; Williamson and FitzPatrick, 2014). This again raises the possibility that the developing eye is robust and can sometimes cope with mutations in genes involved in eye formation.

Genetic robustness is the capacity of organisms to withstand mutations, such that they show little or no phenotype, or compromised viability (Felix and Barkoulas, 2015; Wagner, 2005). This inherent property of biological systems is wired in the genetic and proteomic interactomes and enhances the chance of viability of individuals in the face of mutations. High throughput reverse mutagenesis projects and the emergence of CRISPR/Cas9 gene editing techniques have highlighted the fact that homozygous loss of function mutations in many genes generate viable mutants with no overt phenotype (Varshney et al., 2015; Dickinson et al., 2016; Meehan et al., 2017). Across phyla, mutations in single genes are more likely to give rise to viable organisms than to show overt or lethal phenotypes. For instance, it is estimated that zygotic homozygous null mutations in just ∽7% of zebrafish genes compromise viability before 5 days post fertilisation (Kettleborough et al., 2013) and 8-10% between day 5 and 3 months (Shawn Burgess, personal communication); and compromised viability is predicted following loss of function for about 35% of mice genes (Dickinson et al., 2016; Meehan et al., 2017). Furthermore, apparently healthy viable homozygous or compound heterozygous ‘gene knockouts’ have been found for 1171 genes in the Icelandic human population (Sulem et al., 2015) and for 1317 genes in the Pakistani population (Saleheen et al., 2017).

In some cases, the lack of overt phenotype may be due to redundancy in gene function based on functional compensation by paralogous or related genes (Barshir et al., 2018, Hurles, 2004, Wagner, 1996). We can assume that genes that do not express a phenotype when mutated are not lost to genetic drift because in some way they enhance the fitness of the species. For instance, even though two paralogous *Lefty* genes encoding Nodal signalling feedback effectors have been shown to be dispensable for survival, they do make embryonic development robust to signalling noise and perturbation (Rogers et al., 2017).

Genetic compensation for deleterious mutations is a cross-species feature (El-Brolosy and Stanier, 2017), and mRNAs that undergo nonsense-mediated decay due to mutations that lead to premature termination codons can upregulate the expression of paralogous and other related genes (El-Brolosy et al., 2018). However, only a fraction of genes have paralogues and other compensatory mechanisms must contribute to the ability of the embryo to cope with potentially deleterious mutations. One such mechanism is distributed robustness, which can emerge in gene regulatory networks (Wagner, 2005). This kind of robustness relies on the ability of the network to regulate the expression of genes and/or the activity of proteins within the network, such that homeostasis is preserved when one of its components is compromised (Davidson, 2010; Peter and Davidson, 2016).

Maternal-zygotic *tcf7l1a* mutant zebrafish have been previously described as lacking eyes (Kim et al., 2000). In this study, we show that expression of this phenotype is dependent on the genetic background. We find that *tcf7l1a* mutants can develop functional eyes and are viable, and that this is not due to compensatory upregulation of other *lef/tcf* genes. Despite the presence of functional eyes, the eye field in *tcf7l1a* mutants is only half the size of the eye field of wildtype embryos, indicating an early requirement for *tcf7l1a* during eye field specification. We further show that this requirement is cell autonomous, revealing a striking dissociation between the early role and requirement for Tcf71a in eye field specification and the later absence of an overt eye phenotype. Subsequent to compromised eye field specification, *tcf7l1a* mutant eyes recover their size by delaying neurogenesis and prolonging growth in comparison to wildtype eyes. This compensatory ability of the developing eye was also observed when cells are removed from the wild type optic vesicles. All together, our study suggests that the loss of Tcf7l1a does not trigger any genetic compensation or signalling pathway changes that restore eye field specification; instead, the developing optic vesicle shows a remarkable ability to subsequently modulate its development to compensate for the early, severe loss of eye field progenitors.

The penetrance and expressivity of eye phenotypes appears to be dependent on complex genetic and environmental interactions (Gestri et al. 2009; Kaukonen et al., 2018; Prokudin et al., 2014). Thus, we speculated that *Tcf7l1a* mutant eyes may be sensitised to the effects of additional mutations. Here we show this is indeed the case and describe the isolation of three mutations from a recessive synthetic modifier screen in *tcf7l1a* homozygous mutant zebrafish that lead to enhanced/novel eye phenotypes when in combination with loss of *tcf7la* function.

In summary, our work shows that zebrafish eye development is robust to the effects of a mutation in *tcf7l1a* due to growth compensatory mechanisms that may link eye size and neurogenesis. Our study adds to a growing body of research revealing a variety of mechanisms by which the developing embryo copes with the effects of deleterious genetic mutations.

## Results

### The *tcf7l1a*^*m881/m881*^ mutation is fully penetrant but maternal-zygotic mutants show no overt eye phenotype and are viable

The *headless (hdl)*^*m88*^ mutation in *tcf7l1a* (*tcf7l1a*^*-/-*^ from here onwards) was identified because embryos lacking maternal and zygotic (MZ) gene function lacked eyes (Kim et al., 2000). However, no overt defects were observed in zygotic (Z) *tcf7l1a*^*-/-*^ mutants, due to functional redundancy with the paralogous *tcf7l1b* gene (Dorsky et al., 2003). In our facility, *MZtcf7l1a*^*-/-*^ embryos showed a variable eye phenotype, ranging from eyeless, to small and overtly normal eyes, with proportions that varied in clutches from different pairs of fish (not shown). We hypothesised that genetic background effects could be responsible for either enhancing or suppressing the eyeless phenotype. To test this idea, we outcrossed *tcf7l1a*^*-/-*^ fish to *ekkwill* (*EKW*) wildtype fish and identified *tcf7l1*^*+/-*^ carriers by PCR genotyping. After three generations of outcrossing to *EKW* fish, we incrossed *tcf7l1*^*+/-*^ carriers to grow *Ztcf7l1a*^*-/-*^ adults. All *MZtcf7l1a*^*-/-*^ embryos coming from six pairings of Z*tcf7l1a*^*-/-*^ mutant fish developed eyes only slightly reduced in size compared to eyes of wildtype embryos of the same *EKW* strain (100%, n>100; Fig.1A,B).

**Fig.1.**
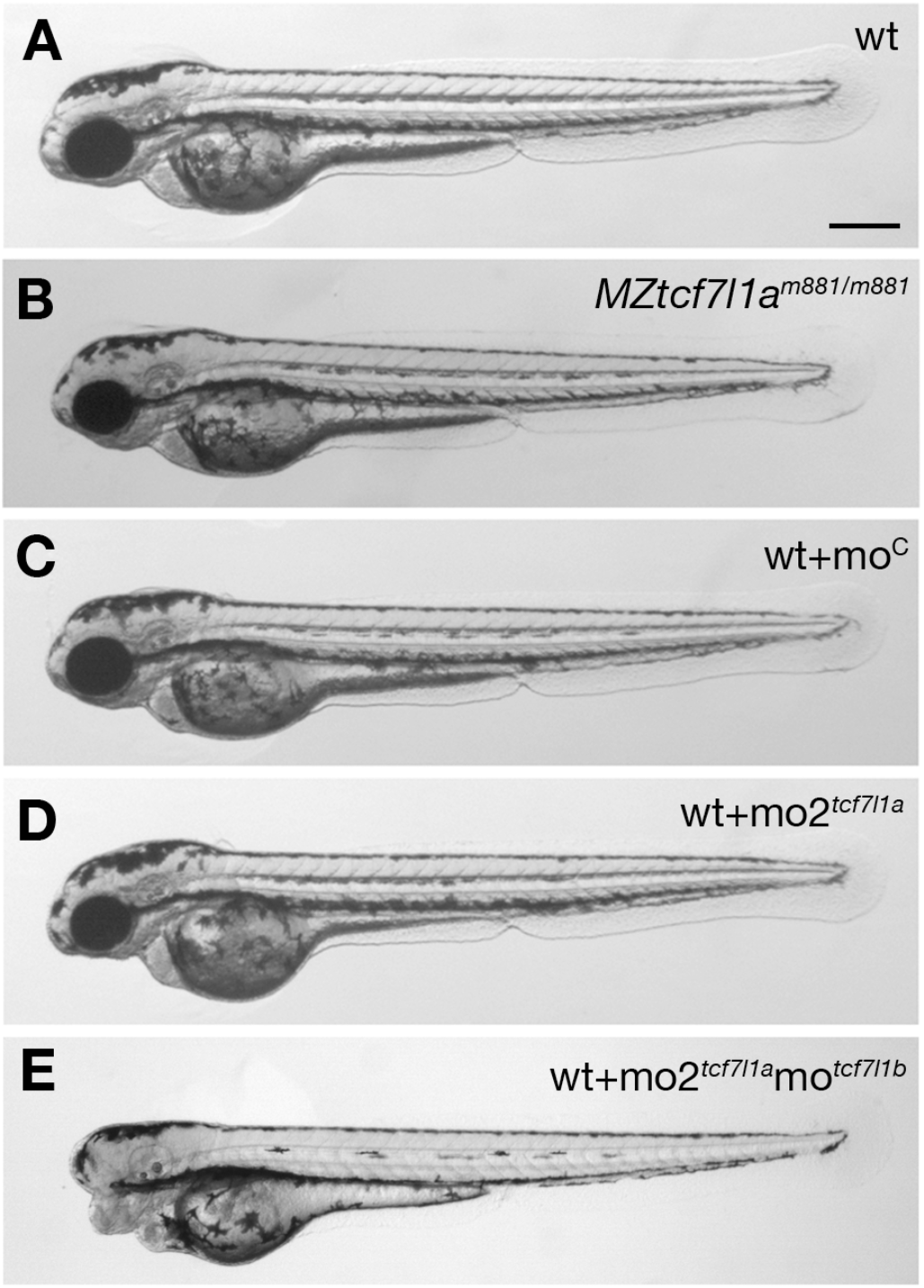
*Tcf7l1a* maternal zygotic (MZ) mutants and morphants have no overt eye phenotype. Lateral views of wildtype (**A**), *MZtcf7l1a*^*-/-*^ (**B**), wildtype injected with control morpholino (**C**), *tcf7l1a* morphant (**D**) and *tcf7l1a/tcf7l1b* double morphant (**E**) embryos at 2 days post fertilisation. Dorsal up, anterior to the left. Scale Bar=250μm.

The *tcf7l1a*^*m881*^ mutation creates a splice acceptor site in intron 7, which leads to a 7 nucleotide insertion in *tcf7l1a* mRNA that gives rise to a truncated protein due to a premature termination codon (Kim et al., 2000). Given that the wildtype splice site in intron 7 is still present in *tcf7l1a* mutants, we assessed whether the lack of phenotype in *MZtcf7l1a*^*-/-*^ mutants could be due to incomplete molecular penetrance as a result of expression of mRNA from both wildtype and mutant splice sites. The chromatogram sequence of the RT-PCR product amplifying exons 7 and 8 in wildtype, mutant and heterozygous embryos shows that only wildtype *tcf7l1a* mRNA is detected in wildtype embryos and only mutant mRNA containing the 7 nucleotide insertion is observed in mutants, while heterozygous embryos produce both wildtype and mutant mRNAs (Fig.S1; Kim et al., 2000). This suggests that the mutant splice site is the only one used in *tcf7l1a*^*-/-*^ embryos. In addition, while overexpression of wildtype *tcf7l1a* mRNA rescues eye formation in embryos in which both *tcf7l1a* and *tcf7l1b* are knocked down, *tcf7l1a*^*m881*^ mutant mRNA does not, confirming that protein arising from the *tcf7l1a*^*m881*^ allele is not functional (not shown; Kim et at., 2000). These observations suggest that the *m881* allele is indeed a null mutation and that *tcf7l1a* is not essential for eye formation.

Supporting a requirement for *tcf7l1a* to form eyes, antisense morpholino knockdown of *tcf7l1a* (mo1^*tcf7l1a*^) leads to eyeless embryos (Dorsky et al., 2003) comparable to the originally described *headless MZtcf7l1a*^*-/-*^ mutant phenotype (Kim et al. 2000). However, the target site for the morpholino used in that study shows considerable sequence homology to the translation start ATG region of other *tcf* gene family members (56-76%; Fig.S2A). This suggests that the mo1^*tcf7l1a*^ phenotype may be due to the morpholino knocking down expression of other *tcf* genes, as has been described for other morpholinos targeting paralogous genes (Kamachi et al., 2008). Indeed, injection of a different *tcf7l1a* morpholino (mo2^*tcf7l1a*^) with low homology to other *tcf* genes (36-45%, Fig.S2B) does not lead to an eyeless phenotype (0.4pMol/embryo, 100%, n>100; Fig.1C,D). *tcf7l1b* morpholino injection on its own shows no overt phenotype (Dorsky et al., 2003) but co-injection of mo2^*tcf7l1a*^ and mo^*tcf7l1b*^ gives rise to eyeless embryos (each at 0.2pMol/embryo, 78.26%, n=92; Fig.1E and see Dorsky et al., 2003). This suggests that even though mo2^*tcf7l1a*^ injection alone results in no phenotype, the morpholino does knockdown *tcf7l1a*.

Together, these results suggest that even though *tcf7l1a*^*-/-*^ is a fully penetrant null mutation, lack of maternal and zygotic *tcf7l1a* function alone does not lead to loss of eyes in all genetic backgrounds.

### *tcf7l1a* loss of function is not compensated by upregulation of other *tcf* genes

Z*tcf7l1a*^*-/-*^ and MZ*tcf7l1a*^*-/-*^ embryos develop eyes whereas embryos lacking both Z*tcf7l1a* and *Ztcf7l1b* do not (Dorsky et al. 2003). Thus, we hypothesised that enhanced expression of the paralogous *tcf7l1b,* or other *lef*/*tcf* genes may compensate for the absence of *tcf7l1a* function, as shown for other mutations (El-Brolosy et al., 2018; Rossi et al., 2015). To test this idea, we assessed the expression of all *lef/tcf* genes by RT-qPCR in sibling wildtype and Z*tcf7l1a*^*-/-*^ mutant embryos at the stage when the eye field has been specified (10 hours post fertilisation; hpf).

Expression levels of *lef*/*tcf* genes did not increase in *Ztcf7l1a*^*-/-*^ mutant embryos, which suggests that there is no compensatory upregulation (Fig.2A, TableS1). As previously shown, *tcf7l1a* undergoes nonsense-mediated decay in mutants resulting in reduced expression levels (Kim et al., 2000; Fig2A; TableS1). *lef1* and *tcf7* levels did not change significantly in mutants and *tcf7l1b* (*tcf3b*) and *tcf7l2* (*tcf4*) expression was actually reduced to 63±6% and 62±8% respectively of wildtype levels (Fig.2A; TableS1). The *otx1b* and *otx2* genes, which are expressed in the anterior neural plate, also showed slightly reduced expression (*otx1b*, reduced to 81±11% and *otx2*, 79±10%) suggesting the anterior neural plate may be slightly reduced in size in mutants. Indeed the domain of the neural plate encompassed by expression of *emx3* around the anterior margin of the neural plate up to the mesencephalic marker *pax2a* (Fig.2 D, E) was reduced to 76% of wildtype size in mutants (n=11, p=0.0041, Fig.2B; TableS2). This indicates that a reduction in the size of the prospective forebrain of *Ztcf7l1a*^*-/-*^ embryos may contribute to the reduced levels of expression of *tcf7l1b, tcf7l2 and otx* genes. Overall these results suggest that *tcf* genes do not show compensatory upregulation in response to loss of *tcf7l1a* function.

**Fig.2.**
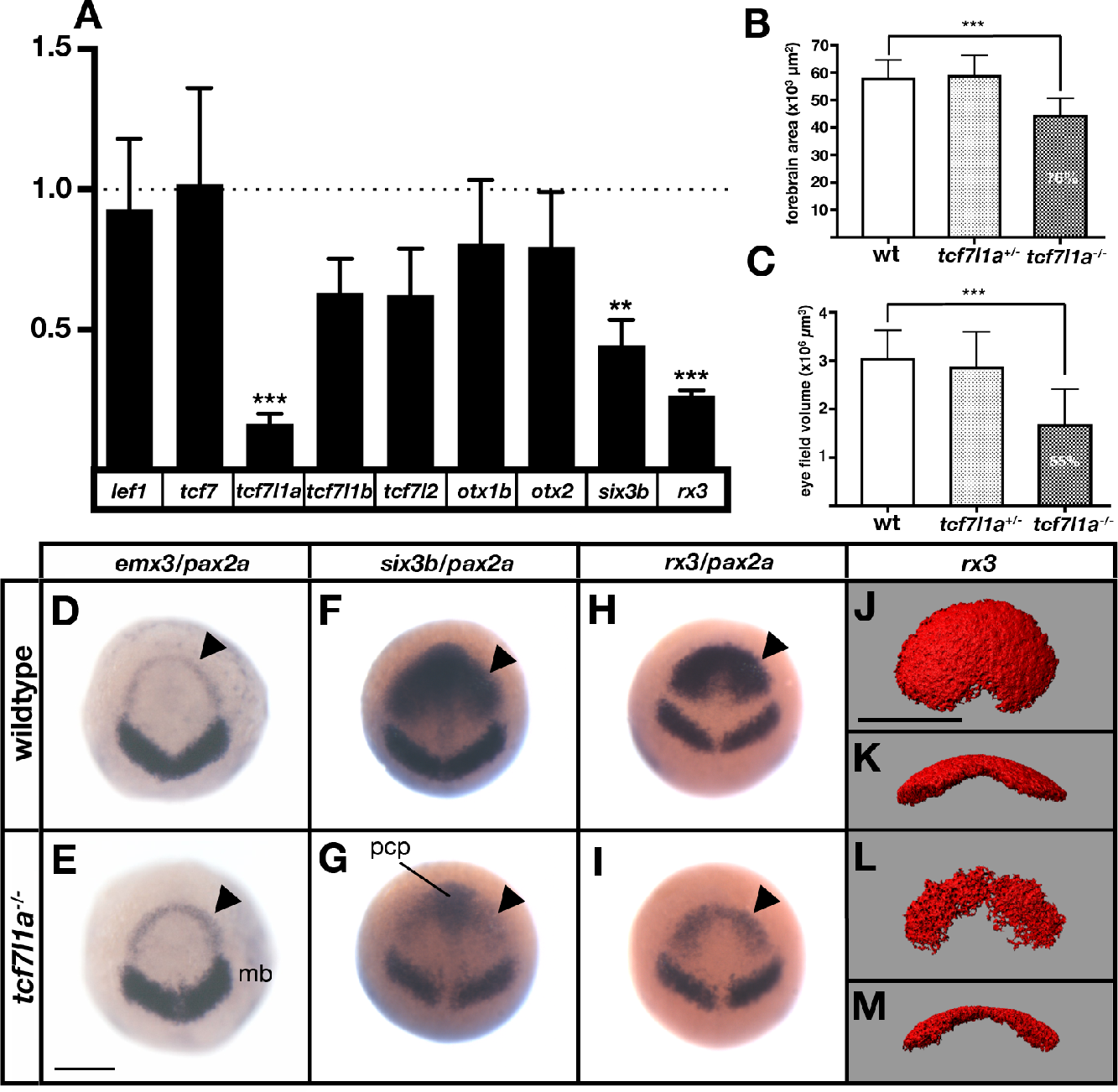
The prospective forebrain and eye field domains of the neural plate are reduced in Z*tcf7l1a*^*-/-*^ mutants. **(A)** Graph showing qRT-PCR quantification of the mRNA levels of *lef1, tcf7, tcf7l1a, tcf7l1b, tcf7l2, otx1b, otx2, six3b* and *rx3* in Z*tcf7l1a*^*-/-*^ mutants relative to wildype embryos at 10hpf. (**B, C**) Quantification of the forebrain domain of the anterior neural plate (**B**) enclosed by *emx3* up to *pax2a* expression by *in situ* hybridisation (data in Table S2), and eye field volume (**C**) by *rx3* fluorescent *in situ* hybridisation confocal volume reconstruction (data in Table S3). (**D-I**) Expression of *emx3* (arrowhead)/*pax2a* (**D, E**), *six3b* (arrowhead)/*pax2a* (**F, G**) and *rx3* (arrowhead)/*pax2a* (**H, I**) in wildtype (**D, F, H**) and Z*tcf7l1a*^*-/-*^ (**E, G, I**) embryos detected by *in situ* hybridisation at 10hpf. (**J-M**) Confocal volume reconstruction of *rx3* fluorescent *in situ* hybridisation in wildtype (**J, K**) and Z*tcf7l1a*^*-/-*^ (**L, M**) mutants at 10hpf. (**J, L**) Dorsal view, anterior to top, and (**K, M**) transverse view from posterior dorsal up. Abbreviations: mb, midbrain; pcp, prechordal plate Scale Bars=250μm.

### Optic vesicles evaginate and form eyes in MZ*tcf7l1a*^*-/-*^ mutants despite a much-reduced eye field

More remarkable than the modest changes in *tcf* and *otx* gene expression was the finding that qRT-PCR showed very reduced expression of eye field genes in Z*tcf7l1a*^*-/-*^ mutant embryos (Fig.2A; *rx3* reduced to 26±1%, p=0.0002 and *six3b* reduced to 44±5%, p=0.0091 of wildtype levels). Consequently, the presence of overtly normal looking eyes in both Z*tcf7l1a*^*-/-*^ and MZ*tcf7l1a*^*-/-*^ embryos is surprising given that *rx3*^*-/-*^ mutant embryos lack eyes due to impaired specification/evagination of the optic vesicles (Loosli et at., 2003; Stigloher et al., 2006). We confirmed that expression of *six3b* and *rx3* is reduced in the anterior neural plate by *in situ* hybridisation in *Ztcf7l1a*^*-/-*^ and *tcf7l1a* morphant embryos (100%,n>40; Fig.2F-I; Fig.S3A,B; similar changes seen in MZ*tcf7l1a*^*-/-*^ mutants, data not shown). The expression of *six3b* is reduced in the eye field but not in the prechordal plate of *Ztcf7l1a*^*-/-*^ mutants, likely explaining why qRT-PCR shows a greater reduction in *rx3* than *six3b* expression (Fig.2F-I; TableS1). Analysis of eye field volume by fluorescent *in situ* hybridisation (FISH) of *rx3* revealed a reduction to 54.7% of wildtype size (n=10, Fig.2C, J-M; TableS3) and intensity of expression within the reduced eye field also appeared reduced (Fig.2H,I).

Further ISH analysis suggests that it is the caudal region of the eye field that is most affected in *Ztcf7l1a*^*-/-*^ mutants. *emx3* expression directly rostral to the eye field is slightly broader in *Ztcf7l1a*^*-/-*^ mutants than wildtypes but expression does not encroach into the reduced eye field (Fig.2D,E; Fig.S4A,B, n=5 each condition). Conversely, expression of the prospective diencephalic marker *barhl2* caudal to the reduced eye field was expanded rostrally at 10hpf (Fig.S4C,D n=5 each condition) and even more evidently at 9hpf (Fig.S4E,F, 13/13 Z*tcf7l1a*^*-/-*^). These observations suggest a caudalisation of the anterior neural plate in Z*tcf7l1a*^-/-^ mutants leading to reduced eye field specification consistent with phenotypes observed in conditions in which Wnt pathway repression is reduced (Heisenberg et al., 2001; Van de Water et al., 2001).

Despite the small size of eye field in *tcf7l1a***^-/-^**mutants, optic vesicles appear to evaginate normally. Time lapse analysis of optic vesicle evagination using the *Tg(rx3:GFP)*^*zf460Tg*^ transgene to visualise eye field cells (Brown et al., 2010) showed that optic vesicle morphogenesis in *Ztcf7l1a*^*-/-*^ embryos proceeds as in heterozygous sibling embryos (Fig.S5A,B; *tcf7l1a*^*+/-*^, n=6 and *Ztcf7l1a*^*-/-*^ n=6; MovieS1 and S2).

### *Tcf7l1a* functions cell-autonomously to promote eye field specification

Although Tcfs regulate the balance between activation and repression of the Wnt/βCatenin pathway during anterior neural plate regionalisation (Kim at el., 2000, Dorski et al., 2003), it is unclear if Tcf function in the eye field is required for cells to adopt retinal fate. To address this, we determined whether Tcf7l1a function is required cell-autonomously during eye formation by transplanting wildtype and MZ*tcf7l1a*^*-/-*^ GFP labelled (GFP+) cells into wildtype and mutant hosts and analysing the expression of *rx3* when eye specification has occurred (100% epiboly; Fig.3).

**Fig.3.**
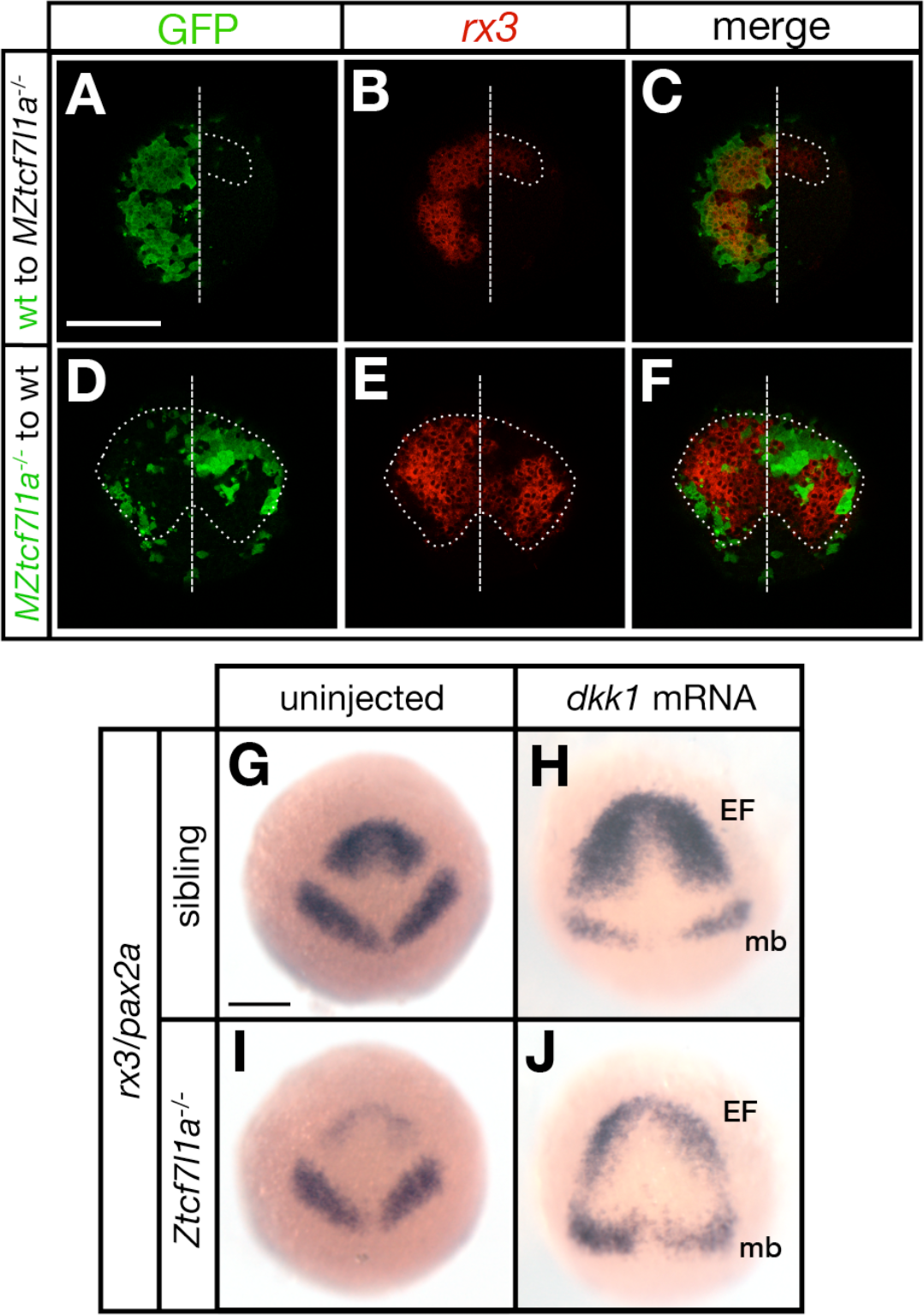
Tcf7l1a cell autonomously promotes *rx3* expression in the eye field. (**A-F**) Dorsal views of confocal images of *rx3* mRNA expression (red) detected by florescent *in situ* hybridisation at 10hpf in the anterior neural plates of chimeric embryos containing transplants of (**A-C**) wildtype (GFP+) donor cells in *MZtcf7l1a*^*-/-*^ host embryos, and (**D-F**) *MZtcf7l1a*^*-/-*^ (GFP+) donor cells in wildtype host embryos. Dotted line outlines eye fields; note in A-C that *rx3* expression extends considerably caudal to the reduced mutant eye field on the side of the neural plate containing wild-type cells. Dashed line marks the embryo midline. **(G-J)** *In situ* hybridisation of *rx3* and *pax2a* in sibling (**G, H**) and *Ztcf7l1a*^*-/-*^ (**I, J**) embryos, uninjected (**G, I**) or injected with 50pg of *dkk1* mRNA (**H, J**) at 9hpf. Abbreviations; EF, eyefield; mb, midbrain Scale Bars=200μm.

Transplants of wildtype cells to MZ*tcf7l1a*^-/-^ mutant embryos led to the recovery of *rx3* expression exclusively restricted to the wildtype GFP+ cell clones (13/13 transplants, Fig.3A-C). However, the border of the GFP+ wildtype clones showed less *rx3* expression, suggesting that cells at the edge of the graft are subject to cell non-autonomous signalling effects from cells surrounding the clone. Conversely MZ*tcf7l1a*^-/-^ mutant GFP+ cells express much lower levels of *rx3* than wild type neighbours when positioned in the eye field of wildtype embryos (9/9 transplants, Fig.3D-G). The reduction in *rx3* expression was limited to the MZ*tcf7l1a*^-/-^GFP+ mutant cells. Control transplants of cells from wildtype donor embryos to wildtype hosts showed no effect on *rx3* expression (not shown). Consistent with a cell autonomous role for Tcf7l1a in eye formation, overexpression of the Wnt inhibitor Dkk1 (Hashimoto et al. 2000) expanded the anterior neural plate in both wildtype and *tcf7l1a* morphants, but *rx3* expression and eye field size remained much smaller in the enlarged anterior plate (Fig.3H-K)).

All together, these results support a cell-autonomous role for Tcf7l1a in promoting eye field specification.

### Eye size in *Ztcf7l1a*^*-/-*^ embryos recovers with growth kinetics similar to wildtype embryos

Despite a much-reduced eye field, eyes in Z*tcf7l1a*^*-/-*^ fry and adults seem indistinguishable from those in wildtype siblings. Indeed, optokinetic responses of Z*tcf7l1a*^*-/-*^ and wildtype 5dpf larvae showed no significant differences at any of the four tested spatial frequencies (Fig.S6), suggesting that by this stage, Z*tcf7l1a*^*-/-*^ eyes are functional and have a visual acuity comparable to that of wildtype siblings. Consequently, although Z*tcf7l1a*^*-/-*^ embryos show a robust and severe neural plate patterning phenotype, this recovers over time. To explore how this recovery happens, we measured the eye size in Z*tcf7l1a*^*-/-*^ embryos from 24 to 96hpf (Fig.4A, C-L; TableS4).

**Fig.4.**
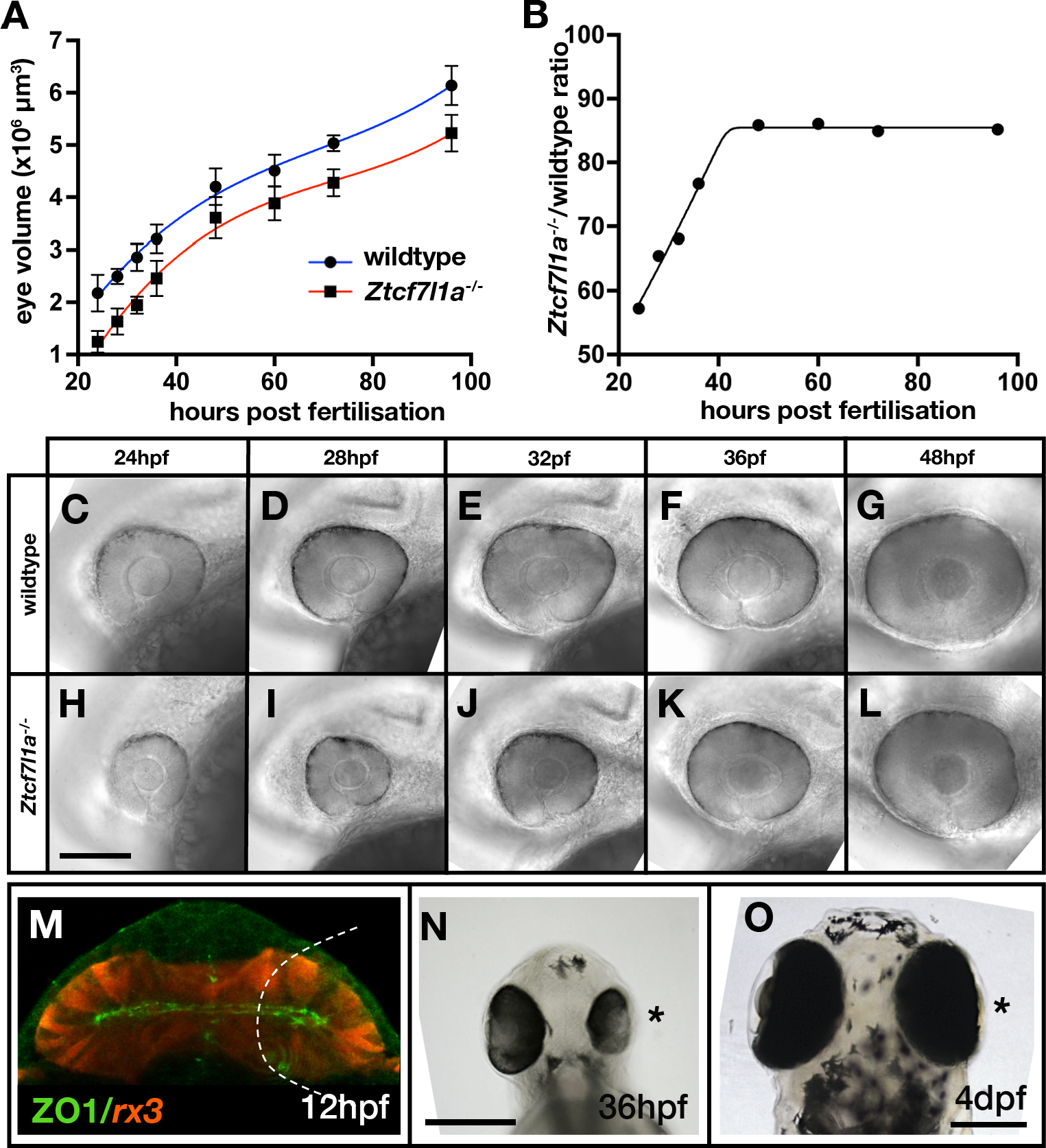
Eye size recovers in Z*tcf7l1a*^*-/-*^ mutant and eye vesicle-ablated embryos. (**A**) Growth kinetics of the eye in wildtype (blue line) and *Ztcf7l1a*^*-/-*^ (red line) embryos at stages indicated (data in Table S4). **B**) Plot showing the ratio of *Ztcf7l1a*^*-/-*^ to wildtype eye volume from data in (**A**). (**C-L**) Lateral views (dorsal up, anterior to left) of wildtype (**C-G**) and *Ztcf7l1a*^*-/-*^ (**H-L**) eyes at stages indicated above panels. (**M-O**) Eye development following partial ablation of the optic vesicle in wildtype embryos at 5 somite stage. (**M**) Coronal confocal section of evaginating optic vesicles (red) in a wildtype *Tg(rx3:RFP)* 5 somite stage embryo. Dashed line indicates the approximate extent of ablations performed. Eye vesicle-ablated embryo at 36hpf (**N**) and 4dpf (**O**). Asterisk indicates the eye that develops from the ablated optic vesicle. ZO1, zona ocludens 1. Scale bars =200μm.

We estimated eye volumes from retinal profiles (see methods) in Z*tcf7l1a*^*-/-*^ and wildtype embryos. At 24hpf, eye volume in mutants was about 57% of the estimated volume of wildtype eyes at the same stage (Fig.4A,C,H; TableS4). However, by 48hpf mutant eyes are about 85% of the size of eyes in wildtype/heterozygous siblings (Fig.4A, G, L; TableS4). Z*tcf7l1a*^*-/-*^ eye size does not further recover beyond this time point and up to 5dpf (Fig.4B). Eye growth in both wildtypes and Z*tcf7l1a*^*-/-*^ mutants show similar growth kinetics (Fig.4A). This suggests that even though Z*tcf7l1a*^*-/-*^ eyes are smaller, they follow a comparable developmental time-course as wildtype eyes in the early growth phase between 24 and 36hpf but with about 8 hours delay (for instance, a 32hpf Z*tcf7l1a*^*-/-*^ eye is about the same size as a wild-type 24hpf eye).

The temporal shift in eye growth in Z*tcf7l1a*^*-/-*^ mutants is not explained by an overall developmental delay as the position of the posterior lateral line primordium (pLLP) was similar to wildtype at all stages tested (Fig.S7).

### Eye size recovers after physical ablation of much of the optic vesicle

To assess if the size recovery is a general feature of eye development, we physically ablated optic vesicle cells in wildtype embryos and assessed the effect on eye growth. Cells were aspirated from one of the two nascent optic vesicles at 12hpf (6 somite stage), leaving approximately the medial half of the vesicle intact (Fig.4M). At 36hpf there was still a clear size difference between the experimental and control eyes (Fig.4N). However, by 4dpf we observed no obvious size difference between control and experimental eyes (n=13/13, Fig.4O). Consequently, the forming eye can effectively recover from either genetic or physical reduction in the size of the eye field/evaginating optic vesicle.

### Neurogenesis is delayed in *tcf71a* mutant eyes

The observation that wildtype and Z*tcf7l1a*^*-/-*^ mutant eyes display similar, but temporally offset, growth kinetics led us to speculate that that retinal neurogenesis might be delayed in Z*tcf7l1a*^*-/-*^ eyes to extend the period of proliferative growth prior to retinal precursors undergoing neurogenic divisions.

In the zebrafish eye, neurogenesis can be visualised by tracking expression of *atoh7* (*ath5*) in retinal neurons starting in the ventronasal retina at ∽28hpf and spreading clockwise across the central retina until it reaches the ventrotemporal side (Masai et al., 2000; Hu and Easter, 1999; Fig.5A-E). Although *atoh7* is induced at a similar time in Z*tcf7l1a*^*-/-*^ as in wildtype eyes, the subsequent progression of expression is delayed (Fig.5F-J, Q; TableS5). Classifying the expression of *atoh7* in 6 categories according to its progression across the neural retina (see legend to Fig.5) revealed that *atoh7* expression in mutant retinas is slow to spread and remains restricted to the ventro-nasal or nasal retina for longer (Fig.5Q). Indeed, between 36 and 40hpf, Z*tcf7l1a*^*-/-*^ retinas express *atoh7* exclusively in the nasal half of the retina (Fig.5H, Q), a phenotype we did not see at any stage in sibling embryo eyes. These data indicate that progression of *atoh7* expression and neurogenesis is delayed by about 8-12 hours in Z*tcf7l1a*^*-/-*^ retinas compared to siblings, a timeframe comparable to the delays seen in optic vesicle growth. In line with our results in Z*tcf7l1a*^*-/-*^ embryos, eye vesicle ablated wildtype retinas also showed delayed neurogenesis compared to control non-ablated contralateral eyes at 36hpf (Fig.5K, L; 3/3 ablated eyes).

Our results suggest that retinal precursors in Z*tcf7l1a*^*-/-*^ eyes remain proliferative at stages when precursors in wildtype eyes are already producing neurons.

**Fig.5.**
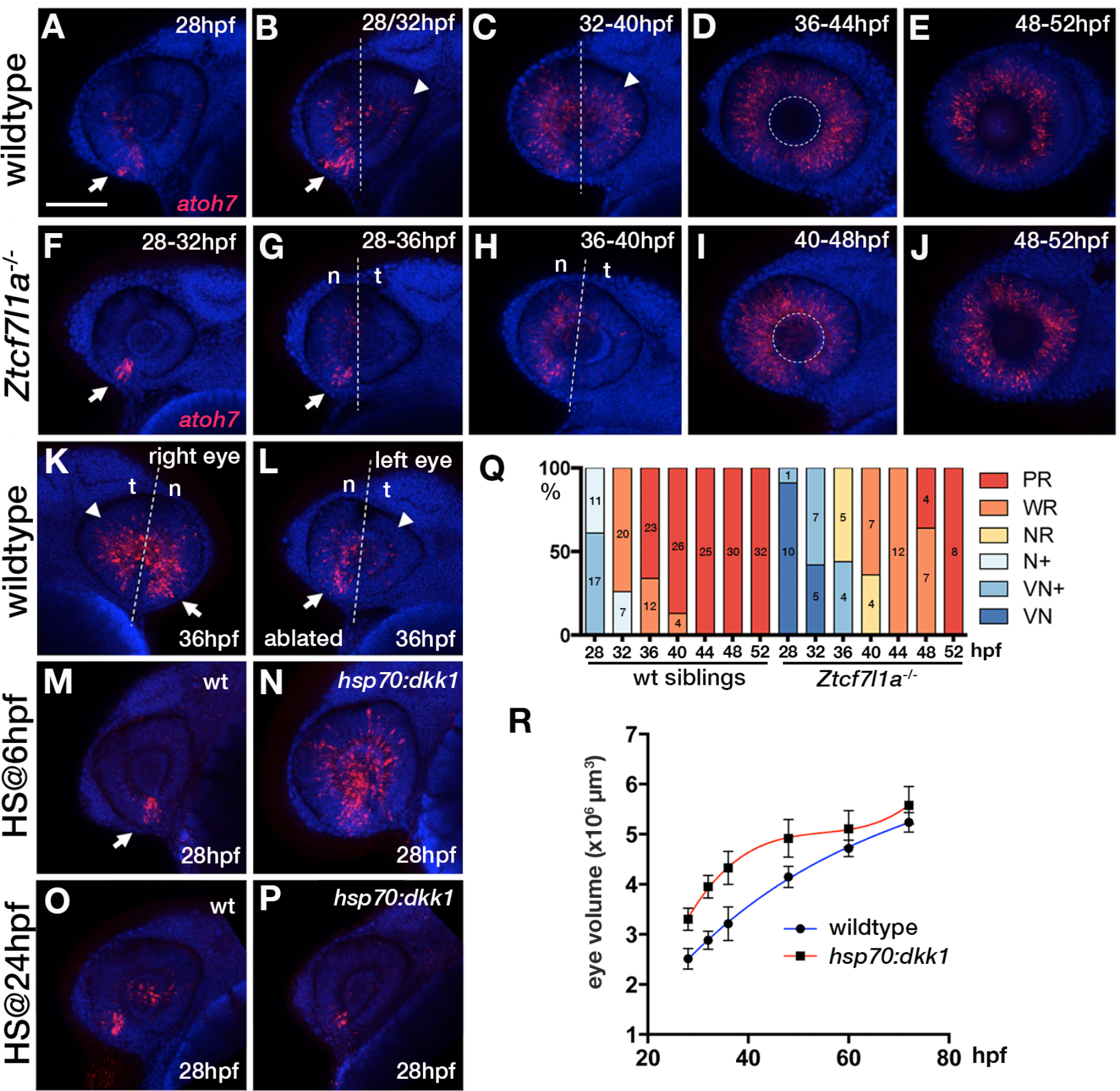
Neurogenesis is delayed in small *tcf7l1a*^-/-^ eyes and accelerated in large eyes following *hsp70:dkk1* overexpression. (**A-P**) Lateral views of eyes showing *atoh7* fluorescent *in situ* hybridisation in wild-type (**A-E, M, O**), *Ztcf7l1a*^*-/-*^ (**F-J**), wildtype left-side optic vesicle-ablated (**K, L**) and *Tg(HS:dkk1)*^*w32*^ (**N, P**) embryos at stages indicated. **M-P)** Wildtype (**M, N**) and heterozygous sibling *Tg(HS:dkk1)*^*w32*^ embryos (**N, P**) heat-shocked at 6hpf (**N, O**) or 24hpf (**P, Q**) for 45’ at 37°C and grown to 28hpf. Anterior is to the left except in (**K**) in which anterior is to the right. Arrows indicate ventro-nasal retina; arrowheads indicate dorso-temporal retina; dashed line approximate the nasal-temporal division; dashed circle marks lens position. Abbreviations: n, nasal, t, temporal. Scale bar =100μm (**Q**) Histogram showing the spatial distribution of *atoh7* expression in sibling and *Ztcf7l1a*^*-/-*^ retinas at the indicated hours post fertilisation (data in Table S5). VN, ventro nasal expression; VN+, ventro-nasal expression plus a few scattered cells; N+, nasal expression plus scattered cells covering the whole retina; NR, nasal retina expression; WR, whole retina expression; PR, expression localised to the peripheral retina. Numbers in bars represent the number of embryos scored for the particular category of *atoh7* expression. (**R**) Plot showing the growth kinetics of the eye in wildtype (blue line) and *Tg(HS:dkk1)*^*w32*^ (red line) embryos at times indicated (data in Table S6).

### Larger eyes undergo premature neurogenesis

Our results are consistent with the idea that neurogenesis may be triggered when the optic vesicle reaches a critical size. To explore this possibility, we generated embryos with larger optic vesicles by overexpressing the Wnt antagonist Dkk1 (Hashimoto et al., 2000). Heatshocking *tg(hsp70:dkk1-GFP)*^*w32*^ transgenic fish at 6hpf led to embryos with eyes ∽34% bigger than control heat shocked embryos by 28hpf (Fig.5M, N, R, n=12; TableS6). After 36hpf, wildtype eyes gradually caught up in size as growth slowed in eyes in *dkk1*-overexpressed embryos (Fig.5R; TableS6).

Neurogenesis was prematurely triggered by 28hpf in the eyes of *dkk1* overexpressing embryos, with many more cells expressing *atoh7* compared to eyes in heat-shocked control embryos (Fig.5M, N, n=7 out of 9 embryos). This result is unlikely to be due to a direct effect of *dkk1* overexpression on neurogenesis as premature neurogenesis is not triggered in *tg(hsp70:dkk1-GFP)*^*w32*^ retinas heat-shocked at 24hpf (Fig.5O, P, n=10, 100%). These results further support a link between eye size and the onset of neurogenesis and the size self-regulating ability of the forming eye.

### ENU modifier mutagenesis screen in *tcf7l1a* mutant background reveals two groups of genetic modifiers

Although eye formation can recover in *tcf7l1a*^-/-^ mutants despite a much smaller eye field, we speculated that eye development in these embryos might be sensitised to showing the effects of additional mutations. To test this, we performed an ENU mutagenesis screen on fish carrying the *tcf7l1a* mutation (Fig.6A).

**Fig.6.**
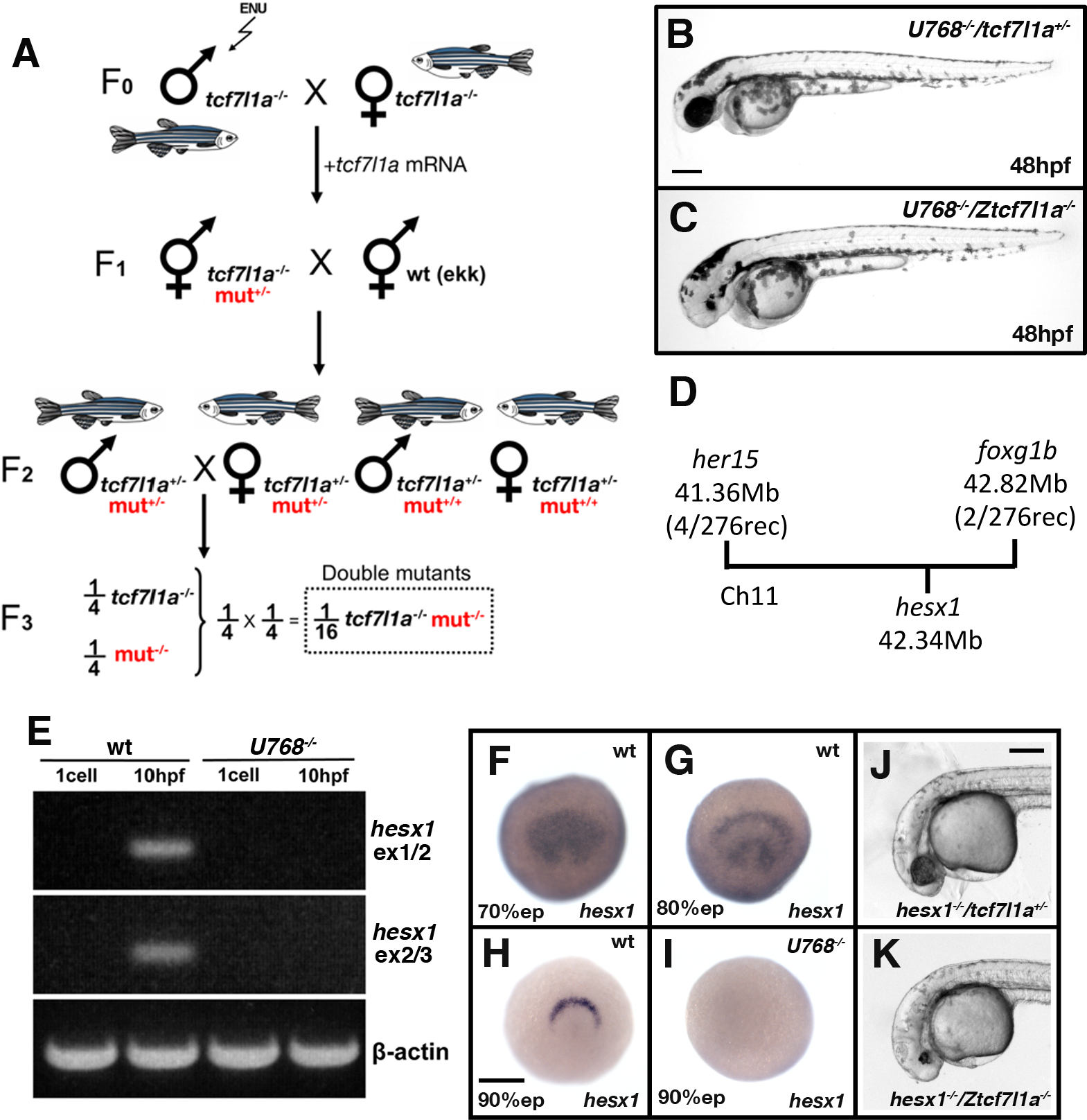
Z*tcf7l1a*^*-/-*^ mutants lacking Hesx1 function fail to form eyes. (**A**) Schematic of the genetic strategy to isolate mutations that modify the *tcf7l1a*^-/-^ mutant phenotype. (**B-C**) *U768* modifier of the *tcf7l1a*^*-/-*^ mutant phenotype. Lateral views of homozygous *U768* embryos that are heterozygous (**B**) or homozygous (**C**) for the *tcf7l1a* mutation. (**D**) Representation of SSLP segregation linkage analysis mapping of E3 modifier of *tcf71a* to a 1.46 megabase (Mb) interval on chromosome 11 (Ch11; rec, recombinants). (**E**) RT-PCR for *hesx1* spanning exons 1-2 (top panel), exons 2-3 (middle panel) and β-actin (bottom panel) on wildtype (lanes 1 and 2) and *U768*^*-/-*^ (lanes 2 and 3) embryo cDNA from 1 cell stage (lanes 1 and 3) and 10hpf (lanes 2 and 4). (**F-I**) *hesx1* in situ hybridisation on wildtype (**F-H**) and *U768*^*-/-*^ (**I**) embryos at epiboly (ep) stages indicated. Dorsal views, anterior up. (**J, K**) Lateral views of *hesx1*^*-/-*^ (Δex1/2) / *tcf7l1a*^*+/-*^ (**J**) and *hesx1*^*-/-*^ (Δex1/2) / *Ztcf7l1a*^*-/-*^ (**K**) embryos. Scale bars=200μm.

Homozygous Ztcf7l1a mutant adult male fish (F_0_ founders) were exposed to four rounds of ENU (van Eeden et al., 1999) and then crossed with Z *tcf7l1a*^-/-^ adult females to generate F_1_ families (Fig.6A). However, possibly because of cellular stress or the synergistic cumulative effect of many mutations induced by ENU, we observed many eyeless F_1_ embryos. To circumvent this problem, we injected 10pg/embryo of zebrafish *tcf7l1a* mRNA to rescue any Tcf-dependent eyeless phenotypes in the F_1_ embryos (Fig.6A). Adult F_1_ fish were outcrossed to EKW wildtype strain. All F_2_ fish were *tcf7l1a*^*+/-*^ and half carried unknown mutations (m) in heterozygosity (Fig.6A). To screen, we randomly crossed F_2_ pairs from each family aiming for at least 6 clutches of over 100 embryos. The probability of finding double Z*tcf7l1a*^*-/-*^*/m*^*-/-*^ embryos for independently segregating mutations is 1/16, hence we would expect to find ∽6 double mutants in 100 embryos. Here, we describe examples of synthetic lethal mutations that lead to microphthalmia/anophthalmia (*U768;* Fig.6C) or eyes that fail to grow (*U762, U901*; Fig.7, 8).

**Fig.7.**
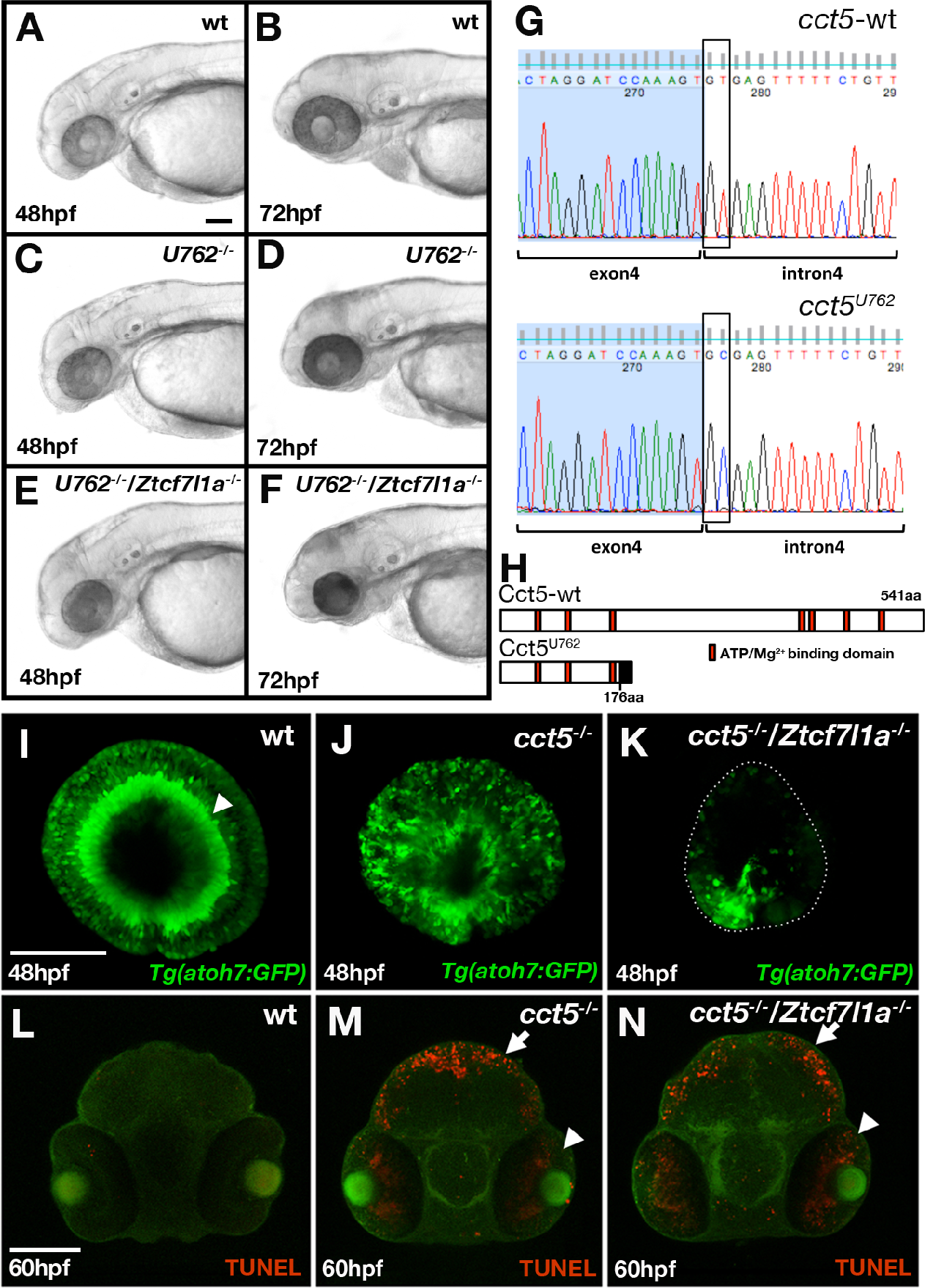
Loss of *tcf7l1a* modifies the *cct5*^*u762/u762*^ mutant eye phenotype. (**A-F**) Laterals view of wildtype (**A, B**), *U762*^*-/-*^ mutants (**C, D**) and double *U762*^*-/-*^*/Ztcf7l1a*^*-/-*^ mutants (**E, F**) at stages indicated. (**G**) DNA sequencing chromatograms of the genomic fragment encompassing the 3’ end of *cct5* exon 4 and 5’ end of intron 4 from wildtype (top) and *cct5*^*U762/762*^ (bottom) embryos. Boxes show the splice donor nucleotides in intron 4. (**H**) Cartoon of wildtype and Cct5^U762^ protein product. Red boxes show Mg^2+^/ATP binding domains, black box indicates nonsense mutant protein stretch. (**I-N**) Anti-GFP immunocytochemistry (green) and TUNEL-labelled apoptotic cells (red) in wildtype (**I, L**), *cct5*^*-/-*^ (*cct5*^*U762/U762*^) (**J, M**) and *cct5*^*-/-*^*/Ztcf7l1a*^*-/-*^ (**K, N**) embryos at 48hpf (**I-K**) and 60hpf (**L-N**). Dotted line in (K) outlines the eye profile. Arrowheads and arrows point to apoptotic cells in the eye and tectum respectively. Scale bar=100μm

### A deletion in the *hesx1* locus is a modifier of the *tcf7l1a*^*-/-*^ phenotype that leads to loss of eyes

*tcf7l1a*^*-/-*^ embryos that are homozygous for the *U768* mutation are eyeless (Fig.6B, C) whereas homozygous *U768* mutants with one or no mutant *tcf7l1a* alleles show no eye phenotype. *U768* was mapped by SSLP segregation analysis (Kelly et al., 2000) to a 1.46Mb interval between 41.36Mb (4 recombinants/274meioses) and 42.82Mb (2 recombinants/274meioses) on chromosome 11 in GRCz10 assembly (Fig.6D; TableS7). Within this interval is *hesx1*, which morpholino knock-down experiments had previously suggested to genetically interact with *tcf7l1a* (Andoniadou et al., 2011). Primers for *hesx1* cDNA failed to amplify in *U768/*Z*tcf7l1a*^*-/-*^ eyeless embryo cDNA samples. Using a primer set that spans the *hesx1* locus, we found that all *U768/Ztcf7l1a*^*-/-*^ eyeless embryos have a ∽2700bp deletion that covers *hesx1* exons 1 and 2 *(hesx1*^*Δex1/2*^; Fig.S8); this was unexpected as deletions are not normally induced by ENU (see below). Sequencing of the *hesx1* locus reveals that there is a polyA stretch of approximately 80 nucleotides followed by a 33 AT microsatellite repeat on the 3’ end of intron 2 that may have generated a chromosomal instability that led to the deletion of exons 1 and 2 (Fig.S8). As a consequence of the deletion, *hesx1* mRNA was not detected by RT-PCR or *in situ* hybridisation in *U768* homozygous embryos (Fig.6E, H, I). We further confirmed that only U768-F_2_ embryos that are homozygous for both the *tcf7l1a* mutation and *hesx1*^*Δex1/2*^ are eyeless (Fig.6B,C).

As ENU usually generates point mutations, we speculated that the deletion in *hesx1*^*Δex1/2*^ was not caused by our mutagenesis but was already present in one or more fish used to generate the mutant lines. Indeed, we found the same deletion in wildtype fish not used in the mutagenesis project. To confirm that the eyeless phenotype in *U768*/Z*tcf7l1a*^*-/-*^ double mutants is not caused by another mutation induced by ENU, we crossed *Ztcf7l1a*^*-/-*^ fish to one such wildtype *TL* fish carrying *hesx1*^*Δex1/2*^. Incrossing of *hesx1*^*Δex1/2/Δex1/2*^*/tcf7l1a*^*+/-*^ adult fish led to embryos with a very small rudiment of eye pigment with no detectable lens (Fig.6J, K). Genotyping of eyeless and sibling embryos confirmed that only double homozygosity for *hesx1*^*Δex1/2*^*/*Z*tcf7l1a*^*-/-*^ led to the eyeless embryo phenotype (TableS8).

The interaction between *hesx1* and *tcf71a* mutations strikingly illustrates how the developing eye can fully cope with loss of function of either gene alone but fails to form in absence of both gene activities. Additional eyeless families that do not carry the *hesx1* deletion were identified but they remain to be validated and mutations cloned.

### The *tcf7l1a* mutation can enhance the phenotypic severity of mutants with small eyes

*U762* mutants, wildtype for *tcf7l1a* or heterozygous for the *tcf7l1a* mutation, show reduced eye size by 72hpf and this phenotype is considerably more severe in embryos homozygous for the *tcf7l1a* mutation (Fig.7A-F). The *U762* mutation was mapped by SSLP segregation analysis to a 1.69Mb interval between 15.50Mb and 17.19Mb on chromosome 24 (Fig.S9A). Through sequencing candidate genes in the interval (Fig.S9A; TableS9), we identified a mutation in the splice donor of *cct5* (chaperonin containing TCP-1 epsilon) intron 4 (GT>GC, Fig.7G). The mutation leads to the usage of an alternative splice donor in the 3’ most end of *cct5* exon 4, which induces a two nucleotide deletion in the mRNA (Fig.S9B). This deletion changes the reading frame of the protein C-terminal to amino acid 176, encoding a 29aa nonsense stretch followed by a stop codon (Fig.7H; Fig.S9B). The mutation also induces nonsense-mediated decay of the mRNA (not shown). *U762* and *cct5*^*hi2972bTg*^ mutations failed to complement (not shown) supporting the conclusion that the mutation in *U762* responsible for the *tcf7l1a* modifier phenotype is in *cct5*. Cct5 is one of the eight subunits of the chaperonin TRiC/TCP-1 protein chaperone complex, which assists the folding of actin, tubulin and many proteins involved in cell cycle regulation (Sternlicht et al., 1993; Dekker et al., 2008; Yam et al., 2008).

As Cct5 is implicated in the folding of cell cycle related proteins, and impairment of the cell cycle could affect neurogenesis in the eye, we assessed retinal neurogenesis by tracking GFP expression in the eyes of *cct5*^*U762/U762*^*/Tg(atoh7:GFP)*^*rw021Tg*^ fish (Fig.7I-K). By 48hpf, wildtype eyes show strong GFP expression in the retinal ganglion cell layer (Fig.7I, n=4, arrow head; Masai et al., 2003) whereas although *cct5*^*U762/U762*^ eye cells do express GFP, lamination of neurons is lost (Fig.7J, n=5). In *cct5*^*U762/U762*^*/*Z*tcf7l1a*^*-/-*^ eyes, GFP-expressing neurons are almost completely restricted to the ventro-nasal retina (Fig.7K, n=6). Subsequently *cct5*^*U762/U762*^*/*Z*tcf7l1a*^*-/-*^ mutants (Fig.7N, n=4, arrowhead) show increased TUNEL labelling of apoptotic cells in the eye compared to single *cct5*^*U762/U762*^ mutants (Fig.7M, arrowhead, n=8).

Homozygous *U901* mutants show a slightly smaller and misshapen eye; this mutation was mapped to *gdf6a* (Valdivia et al., 2016). Unlike Z*tcf7l1a*^*-/-*^ mutants in which eye size recovers, eyes in *gdf6a*^*U901/U901*^*/*Z*tcf7l1a*^*-/-*^ embryos remain smaller than in single mutants or wildtypes (Fig.8A-E). This suggests that the ability to compensate eye size is compromised in absence of both *gdf6a* and *tcf7l1a* function.

**Fig.8.**
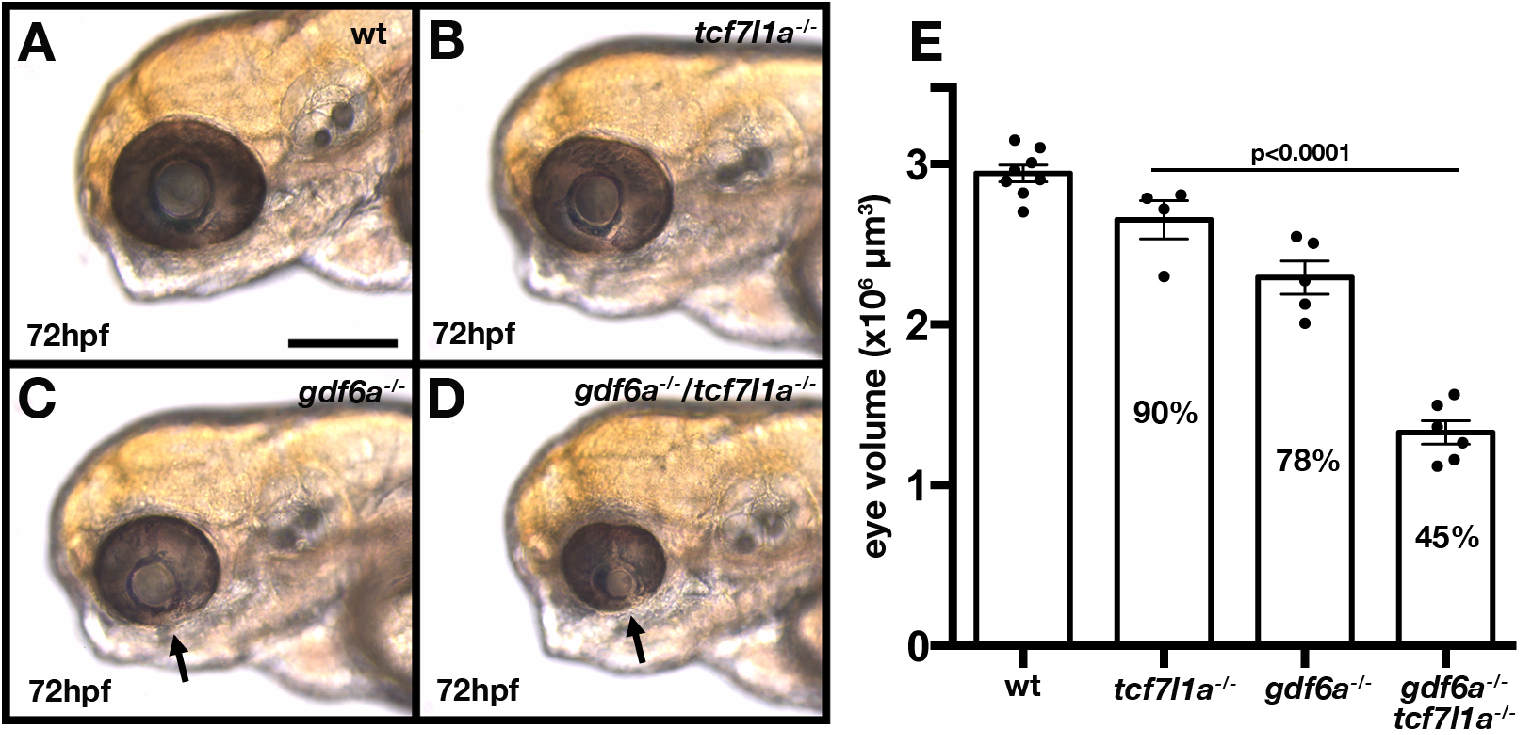
Loss of *tcf7l1a* modifies the *gdf6a*^*U901/U901*^ mutant eye phenotype. (**A-D**) Lateral views of eyes in wildtype (**A**), *Ztcf7l1a*^*-/-*^ (**B**), *gdf6a*^***U901****/****U901***^(**C**) and double *gdf6a****U901****/****U901****/Ztcf7l1a*^*-/-*^ (**D**) embryos at 72hpf. Dorsal up, anterior to left. Arrows indicate the lens. Scale bar=200μm. (**E**) Eye volume quantification in wildtype, *tcf7l1a*^*-/-*^, *gdf6a*^*-/-*^ and *gdf6a/tcf7l1a* double mutant siblings (data in Table S10). Unpaired t-test.

Altogether, analysis of the interacting mutations reveals that although abrogation of Tcf7l1a function alone has little effect on formation of eyes, it can lead to complete loss of eye formation or more severe eye phenotypes in combination with additional mutations. Consequently, although eye development is sufficiently robust to cope with loss of Tcf71a, mutant embryos are sensitised to the effects of additional mutations.

## Discussion

In this study, we show that although Tcf7l1a is required for cells to adopt eye field identity and express *rx3, tcf7l1a* mutants form normal, functional eyes. This finding reveals a remarkable ability of the developing eye to form normally from an eye field that is half the size of that in the wildtype condition. Tcf function in *tcf7l1a* mutants is not genetically compensated by upregulation of other *tcf* genes nor by other genetic mechanisms that restore neural plate regionalisation and eye field formation. Instead, we find that *tcf7l1a* mutant optic vesicles delay neurogenesis to enable size recovery. We observe a similar effect when optic vesicle cells are physically ablated. In contrast, neurogenesis is prematurely induced in larger optic vesicles, likely depleting progenitors and slowing growth. Our results suggest that size-dependent regulation of the balance between proliferation and differentiation may buffer the developing eye against initial differences in cell number. Although the developing eye can cope with loss of Tcf7l1a function, we speculated that embryos lacking Tcf7l1a would not be robust to the consequences of additional mutations affecting eye formation. In support of this, we identify mutations in three other genes that give synthetically enhanced eye phenotypes when combined with the *tcf7l1a* mutation. This approach facilitates identification of genes that participate in genetic networks that make developing eyes robust to mutations that compromise eye field specification and optic vesicle growth.

### The *tcf7l1a* mutation is fully penetrant with no apparent genetic compensation during neural plate patterning

*Tcf7l1* is a core Wnt pathway transcription factor that can activate or repress genes dependent upon the status of the Wnt signalling cascade (Cadigan and Waterman, 2012). Homozygous *tcf7l1* mutant mice present severe mesodermal and ectodermal patterning defects (Merrill et al., 2004), but the duplication of *tcf7l1* into *tcf7l1a* and *tcf7l1b* in zebrafish has led to functional redundancy (Dorsky et al., 2003).

Although MZ*tcf7l1a* embryos have a severe eye field specification phenotype they still develop normal eyes. We confirmed that the *tcf7l1a*^*m881*^ mutant allele is null, generates no wildtype transcript and that morpholino knock-down specifically of Tcf7l1a does not give an eyeless phenotype. Hence, the originally described MZ*tcf7l1a* eyeless phenotype (Kim et al., 2000) may have been due to genetic background effects modifying the outcome of the *tcf7l1a*^*m881*^ allele. The fact that we were able to recover an eyeless modifier of the *tcf7l1a* phenotype in our own mutagenesis pilot screen lends support to this idea.

At the stage of eye specification, we did not find genetic compensation in *tcf7l1a* mutants by other *tcf* genes. Even though *tcf7l1a* mutants develop eyes, they do so from an eye field that is ∽50% smaller than wild-type. Although we did not find evidence for genetic compensation, and despite *tcf7l1* being duplicated in fish, the fact that neither gene has been lost due to genetic drift suggests that having both genes may confer enhanced fitness and robustness to zebrafish. As an example, paralogous Lefty proteins make Nodal signalling more stable to noise and perturbations during early embryogenesis (Rogers et al., 2017).

We show that Tcf7l1a has a cell-autonomous role in specification of the eye field. Tcf7l1a is required for the expression of *rx3* and consequently is a bona fide eye field gene regulatory network transcription factor that functions upstream to *rx3. tcf7l1a* is expressed very early in the anterior neural plate and so may work alongside *otx, sox, six* and *pax* genes to regionalise the eye-forming region of the neural plate (Beccari et al 2013; Zuber et al. 2003). Considering that it is the repressor activity of Tcf7l1a that promotes eye formation (Kim et al., 2000), the most likely role for Tcf7l1a is to repress transcription of a gene that suppresses eye field formation.

### Compensatory tissue growth confers robustness to eye development

We show that despite the small eye field in *tcf7l1a* mutants, the optic vesicles evaginate and undergo overtly normal morphogenesis. Although *tcf7l1a* mutant eye vesicles are still much smaller than wild-type at 24hpf, we found that their eye growth kinetics are similar. This suggests that the mechanisms that regulate overall growth of the retina in both conditions are comparable albeit delayed in the *tcf7l1a* mutant retina.

Although *atoh7* expression is initiated in the ventronasal retina in *tcf7l1a* mutants at the same stage as in wild-type eyes, the wave of *atoh7* expression that spreads across the retina is delayed by approximately 8-12hrs in mutants. *atoh7* is required for the first wave of neurogenesis in the retina (Brown et al., 2001; Kay et al., 2001; Wang et al., 2001) and thus, the delay we see in *tcf7l1a* mutants suggests that RPCs continue proliferating in mutants at stages when they are already generating neurons in wild-type eyes. We presume that the extended period of proliferative growth due to delayed neurogenesis enables the forming eye to continue growing and recover its size. We observed a similar phenomenon of delayed neurogenesis and prolonged growth when cells were ablated from the optic vesicles. Conversely, *atoh7* spreads precociously in experimentally enlarged optic vesicles. The premature neurogenesis of RPCs in these conditions may contribute to eyes achieving a final size similar to wild-type. All together, our data suggest that the timing of the spread of neurogenesis across the retina is coupled to size of the eye, thereby providing a mechanism to buffer eye size. It is intriguing that the compensatory changes in growth seen in *tcf7l1a* mutant and optic-vesicle ablated eyes seem to occur prior to the establishment of the ciliary marginal zone, which accounts for the vast majority of eye growth (Fischer et al., 2013).

Our results support classical embryology experiments from Ross Harrison, Victor Twitty and others (Harrison, 1929, Twitty and Schwind, 1931, Twitty and Elliott, 1934). These investigators showed that when eye primordia from small-eyed salamander species (*A. punctatum*) were transplanted to larger-eyed salamanders (*A. Tigrinum*) or vice-versa, the eye derived from the grafted tissue formed an eye of a size corresponding to the donor salamander species. Species-specific size differences are also observed in self-organising *in vitro* cultured eye organoids derived from mouse or human embryonic stem cells (Nakano et al., 2012). Our work, together with the experiments in salamanders and organoids, suggests that the developing eye has intrinsic size-determining mechanisms.

Size regulatory mechanisms have been previously described in other species and perhaps most extensively studied in the fly wing imaginal disc (Potter and Xu, 2001). Indeed, many models have been put forward to explain imaginal disk size control (Eder et al., 2017; Irvine and Shariman, 2017; Vollmer et al., 2017). It is evident that the final size of paired structures within individuals is remarkably similar supporting the idea that the mechanisms that control organ/tissue size are mostly intrinsic to the tissue/organ and highly robust.

### Addressing eye robustness through a forward mutagenesis screen on *tcf7l1a* mutant background

Our results indicate that *tcf7l1a* mutant eyes are sensitised to the effects of additional mutations. Indeed, a homozygous deletion of the two first exons of *hesx1* leads to eyeless embryos when in combination with *tcf7l1a*. This result also confirms our previous observations suggesting a genetic interaction between *hesx1* and *tcf7l1a* based upon morpholino knock-down experiments (Andoniadou et al., 2007). Furthermore, both *hesx1* and *tcf7l1a* are expressed in the anterior neural plate including the eye field, and as observed in *tcf71a* zebrafish mutants, *hesx1* mutant mice also show a posteriorised forebrain (Andoniadou et al., 2007; Martinez-Barbera et al., 2000). These and our results suggest that Tcf7l1a and Hesx1 have similar, overlapping functions in the anterior neural plate such that the eyeless phenotype is expressed in zebrafish only when both genes are abrogated. Mutations in *hesx1* lead to anophthalmia, microphthalmia, septo-optic dysplasia (SOD) and pituitary defects in humans and mice (Dattani et al., 1998; Gaston-Massuet et al., 2008; Martinez-Barbera et al., 2000; Thomas et al., 2001). Interaction of *hesx1* mutations with other genetic lesions may also occur in patients carrying Hesx1 mutations, as the phenotypes in these individuals show variable expressivity (McCabe et al., 2011). In these patients, *tcf7l1a* should be considered as a candidate modifier for *hesx1* -related genetic conditions.

*Gdf6a* is a TGFβ pathway member (David and Massagué, 2018) that when mutated in zebrafish results in small mis-patterned eyes, neurogenesis defects and retino-tectal axonal projection errors (Gosse and Baier, 2009; French et al., 2009). In humans, mutations in *GDF6* have been identified in anophthalmic, microphthalmic and colobomatous patients (Asai-Coakwell et al., 2009) as well as in some cases of Leber congenital Amaerurosis (Asai-Coakwell et al., 2013). Double *gdf6a*^*U768*^*/tcf7l1a* mutant eyes are smaller than both single mutants and fail to recover their size at later stages. This suggests that *gdf6a/tcf7l1a* double mutant optic vesicles do not show compensatory growth as *tcf7l1a* mutants. It is intriguing that *gdf6a* mutants show premature expression of *atoh7* and neurogenesis (Valdivia et al. 2016). If this phenotype is epistatic to the tissue size compensatory mechanisms, then this may contribute to the lack of compensatory proliferative growth in double mutant eyes.

Mutations in *cct5* in combination with *tcf7l1a* also led to phenotypes in which reduced eye size failed to recover. *cct5* codes for the epsilon subunit of the TCP-1 Ring Complex (TRiC) chaperonin that is composed of eight different subunits that form a ring, the final complex organised as a stacked ring in a barrel conformation (Yebenes et al., 2011). *In vitro* studies indicate TRiC chaperonin mediates actin and tubulin folding (Sternlicht et al., 1993); however, it also assists in the folding of cell cycle-related and other proteins (Dekker et al., 2008; Yam et al., 2008). A mutation in *cct2* has been found in a family with Leber congenital ameurosis retinal phenotype (Minegishi et al., 2016; Minegishi et al., 2018) and mutations in *cct4* and *cct5* have been related to sensory neuropathy (Pereira et al., 2017; Lee et al., 2003; Hsu et al., 2004; Bouhouche et al., 2006). Similar to our *cct5* mutant, *cct1, cct2, cct3, cct4 and cct8* mutant zebrafish show retinal degeneration (Berger et al., 2018; Matsuda and Mishina, 2004; Minegishi et al., 2018), suggesting that the *cct5* mutant phenotype is due to abrogation of TRiC chaperonin function, and not due to loss of a *cct5* specific role. Double *cct5/tcf7l1a* homozygous mutant eyes degenerate prematurely and to a greater extent than *cct5* single mutants, and neurogenesis is also severely compromised. This shows that the consequence of *cct5* loss of function is exacerbated by the lack of *tcf7l1a* function, although it is currently unclear how such an interaction might occur. However, this genetic interaction does highlight that in some conditions a gene of pleiotropic function, like *cct5,* can lead to a specific phenotype in the eye.

Anophthalmia and microphthalmia are generally associated with eye field specification defects (Reis and Semina, 2015), but given that normal eyes can still develop from a much reduced eye field, further analysis of the genetic and developmental mechanisms that lead to small or absent eyes is warranted. Our isolation and identification of modifiers of *tcf7l1a* highlights the utility of genetic modifier screens to identify candidate genes underlying congenital abnormalities of eye formation. Indeed, given that Tcf7l1a itself can now be classified as a *bona fide* gene in the eye transcription factor regulatory network, it should be considered when screening patients with inherited eye morphological defects.

## Author Contributions

RY and SW conceived the project and analysed the data; RY, FC, TH, GG, EA, AK, JR and IB performed the experiments; RY, FC, TH, HS, QS, LL and CW performed the genetic screen. RY and SW wrote the paper with input from all co-authors but primarily from FC, HS, TH, QS and GG. SW secured the funding of this project.

## Acknowledgements

We are grateful to Ajay Chitnis and Richard Dorsky for sharing reagents and fish lines, colleagues for support and discussions and the UCL fish facility for fish care. This work was generously supported by funding from a Marie Curie Incoming International Fellowship to RY, Wellcome Trust grants to SW and RY, an MRC Programme grant to SW and GG, Royal Society International Joint Project funding to SW and MA and FONDAP (15090007) to MA.

## Methods

### Animal use, mutant and transgene alleles, genotyping and heat shock

Adult zebrafish were kept under standard husbandry conditions and embryos were obtained by natural spawning. Wildtype and mutant embryos were raised at 28.5°C and staged according to Kimmel et al. (1995). To minimise variations in staging, embryos were collected every 30 minutes and kept separate clutches according to their time of fertilisation. Fish lines used were *tcf7l1a/headless(hdl)*^*m881*^ (Kim et al., 2000), *cct5*^*hi2972bTg*^ (Amsterdam et al., 2004), *cct5*^*U762*^, *gdf6a*^*U768*^ (Valdivia et al., 2016), *hesx1*^U910^, *Tg(atoh7:GFP)*^*rw021Tg*^ (Masai et al., 2000), *Tg(hsp70:dkk1-GFP)*^*w32*^ (Stoick-Cooper et al., 2007) and *Tg(rx3:GFP)*^*zf460Tg*^ (Brown et al., 2010). Genomic DNA was isolated by HotSHOT method (Suppl. Materials and methods) and all the alleles except for *cct5*^*hi2972bTg*^ were genotyped by KASP assays (K Biosciences, assay barcodes: 1077647141 (*cct5*^*U762*^), 1077647146 (*gdf6a*^*U768*^), 172195883 (this assay discriminates a SSLP 500bp from the 3’ end of the deletion in *hesx1*^*U901*^), 1145062619 (*tcf7l1a*^*m881*^)) using 1μl of genomic DNA for 8μl of reaction volume PCR as described by K Biosciences.

For heatshock (HS) gene induction, embryos from a heterozygous *Tg(hsp70:dkk1-GFP)*^*w32*^ to wild type cross were moved from embryo media at 28.5°c to 37°C at 6hpf or 20hpf for 45minutes, and then back to 28.5°C embryo media. Three hours post HS, embryos were separated in controls (GFP-) and HS experimental (GFP+) groups, and fixed at the stages described in results.

### ENU mutagenesis and mutant mapping

Homozygous male *tcf7l1a*^*m881*^ fish were exposed to four rounds of ENU according to Van Eeden et al. (1999). Details of the mutagenesis pipeline are in the results section. Embryos from incrosses of carriers of the *cct5*^*U762*^ or *gdf6a*^*U768*^ mutations, which show a phenotype as homozygous embryos independently of mutations in *tcf7l1a*, were identified for the described eye phenotype at 3dpf to avoid ambiguity and false positives. For rough mapping, batches of 30 mutants and 30 siblings were fixed in methanol and genomic DNA was extracted by proteinase K protocol. This gDNA was then used for bulk segregant analysis PCR to test a library of 245 polymorphic SSLP variants spanning the whole zebrafish genome (Stickney et al., 2002). SSLP markers heterozygous in the sibling samples and homozygous in the mutant sample were confirmed on gDNA samples of 12 mutant and 12 sibling individuals. Markers that showed linkage to a locus were tested on additional mutant samples, and more SSLP markers were tested for the mapped region until a genomic interval was defined.

Homozygous *tcf7l1a/hesx1*^*U901*^ mutant carriers were incrossed, and eyeless embryos and siblings were fixed in methanol. Rough mapping was carried out as above but in this case sibling embryos used for bulk segregant analysis were genotyped for *tcf7l1a*^*m881*^ and only homozygous mutants with eyes were included in the sibling pool.

### mRNA synthesis, embryo microinjection and morpholinos

mRNA for overexpression was synthesised using RNA mMessage mMachine transcription kits (Ambion). One to two cell stage embryos were co-injected with 10nl of 5pg of GFP mRNA and morpholinos or *in vitro* synthesised mRNA at the indicated concentrations. Only embryos with an even distribution of GFP fluorescence were used for experiments. Morpholino sequences: mo2 t*cf7l1a* (5’ AGG CAT GTT GGC ACT TTA AAT G 3’), mo^*tcf7l1b*^ (5’-CAT GTT TAA CGT TAC GGG CTT GTC T-3’; Dorsky et al., 2002) and mo^c^ (TGT TGA AAT CAG CGT GTT CAA G). *tcf7l1a*^*m881/m881*^ embryos injected with mo^*tcf7l1b*^ phenocopy the loss of eye phenotype seen in *tcf7l1a*^*m881/m881*^*/tcf7l1b*^*+/zf157tg*^ double mutants (Young and Wilson, unpublished).

### RNA extraction, reverse transcription and qPCR

Total RNA and genomic DNA were isolated from individual embryos at 10hpf following Life Technologies Trizol protocol. cDNA was synthesised by reverse transcription using SuperscriptII (Life Technologies) with 200ng of total RNA to a final volume of 40μl and oligo dT for priming. The cDNA reaction was diluted 10 times and 5μl were used in 25μl final volume reactions using GoTaq qPCR Master mix (Biorad). Each experimental condition was processed in technical and biological triplicates. All primers used had PCR efficiencies within the 90-100% range. Sequences of the primers used are in the supplementary technical materials. Wildtype and *tcf7l1a*^*m881*^ mutant cDNA fragment spanning the *tcf71a* exon 7/8 border for DNA sequencing were amplified with primers P2 and P3 (Kim et al., 2000).

### *In situ* hybridisation and probe synthesis

Whole mount in situ hybridisation was performed using digoxigenin (DIG) and fluorescein (FLU)-labelled RNA probes according to standard protocols (Thisse and Thisse, 2008). Probes were synthesized using T7 or T3 RNA polymerases (Promega) according to manufacturers’ instructions and supplied with DIG or FLU labelled UTP (Roche). Probes were detected with anti-DIG-AP (1:5000, Roche) or anti-FLU-AP (1:10000, Roche) antibodies and developed with NBT/BCIP mix (Roche), for regular microscopy or Fast Red (Sigma) or CY-3 tyramide (cite) substrate for confocal analysis.

### Quantification of eye profile, eye volume, and posterior lateral line primordium (pLLP) position

The sizes of eye profiles were quantified from lateral view images of PFA-fixed embryos by delineating the eye using Adobe Photoshop CS5 magic wand tool and measuring the area of pixels included in the delineated region. The surface area was then transformed from px^2^ to μm^2^. The eye profile and eye volume were calculated from confocal imaging of *vsx2* in situ hybridisation stained embryos at 24hpf. The eye volume/eye profile ratio average from 10 embryos was 53.24. This ratio was used to estimate eye volume from eye profile area and assumes that the profile area to eye volume ratio is constant after 24hpf.

pLLP migration was measured by analysing the position of the posterior end of the primordium relative to the somite boundary labelled by *in situ* hybridisation with *eya1* and *xirp2a* respectively.

### Confocal microscopy and image analysis

Confocal imaging was performed on a Leica TCS SP8 confocal microscope. For time lapse analyses, the stage was set in an air chamber heated to 28.5°C. Live embryos were immobilized in 1% low melting point agarose (Sigma) and 0.016% Tricaine (Sigma) to anesthetize. Image volume analysis measurement was performed on Imaris 7.7.0 and Fuji.

### Cell transplantation

WIldtype or *MZtcf7l1a*^*-/-*^ embryos used as donors were injected with 50pg of GFP mRNA at 1 cell stage. At 3-4hpf, blastula stage, dechorionated donor and host embryos were mounted in 3% methylcellulose in fish water supplemented with 1% v/v penicillin/streptomycin (5,000 units penicillin and 5mg streptomycin per ml) and viewed with a fixed-stage compound microscope (Nikon Optiphot). Approximately 30-40 cells were taken from the animal pole of donors and transplanted to approximately the same position in hosts by suction using an oil-filled manual injector (Sutter Instrument Company). Embryos were moved to 1% penicillin/streptomycin supplemented fish media and fixed at 10hpf.

### Eye vesicle cell removal

Embryos were mounted in 1% low melting point agarose in Ringer’s solution supplemented with 1% v/v penicillin/streptomycin. A slice of set agarose was removed to expose one of the eyes and a drop of mineral oil (sigma) was placed over the target eye to dissolve the epidermis (Picker et al., 2009). After two minutes the oil drop was removed and optic vesicle cells were sucked out with a capillary needle filled with mineral oil. Embryos were left to recover for half an hour before being released from the agarose.

### Optokinetic response

Optokinetic responses were examined using a custom-built rig to track horizontal eye movements (optokinetic nystagmus) in response to whole-field motion stimuli. Larvae at 4 dpf were mounted in 1% low melting point agarose in fish water and analysed at 5 dpf. The agarose surrounding the eyes was removed to allow normal eye movements. Sinusoidal gratings with spatial frequencies of 0.05, 0.1, 0.13 and 0.16 cycles/degree were presented on a cylindrical diffusive screen 25 mm from the centre of the fish’s head with a MicroVision SHOWWX+ projector. Gratings had a constant velocity of 10 degrees/s and changed direction and/or spatial frequency every 20 s. Eye movements were tracked under infrared illumination (720 nm) at 60 Hz using a Flea3 USB machine vision camera and custom-written software. Custom-designed Matlab code was used to extract the eye velocity (degrees per second).

---

**Fig S1.**
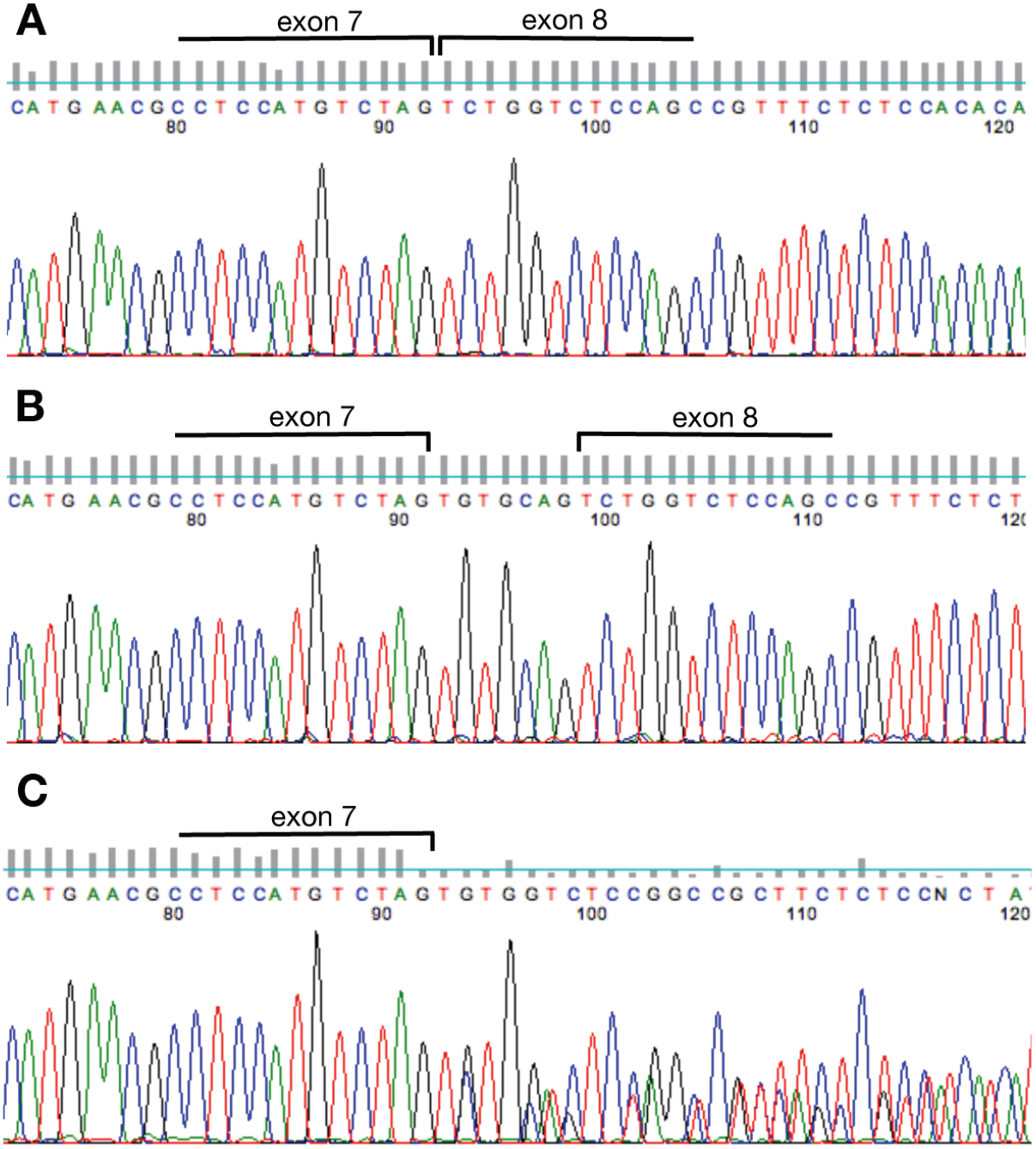
Sequence of the *tcf7l1a* exon7/8 boundary. RT-PCR chromatogram sequence of *tcf7l1a* exon7/8 fragment showing the expected intron splice in wildtype embryos (**A**). *tcf7l1a* **^*-/-*^**mutants show an unambiguous inclusion of 7 nucleotides from intron 7 (**B**) and a mixed read in mRNA coming from heterozygous siblings (**C**).

**Fig S2.**
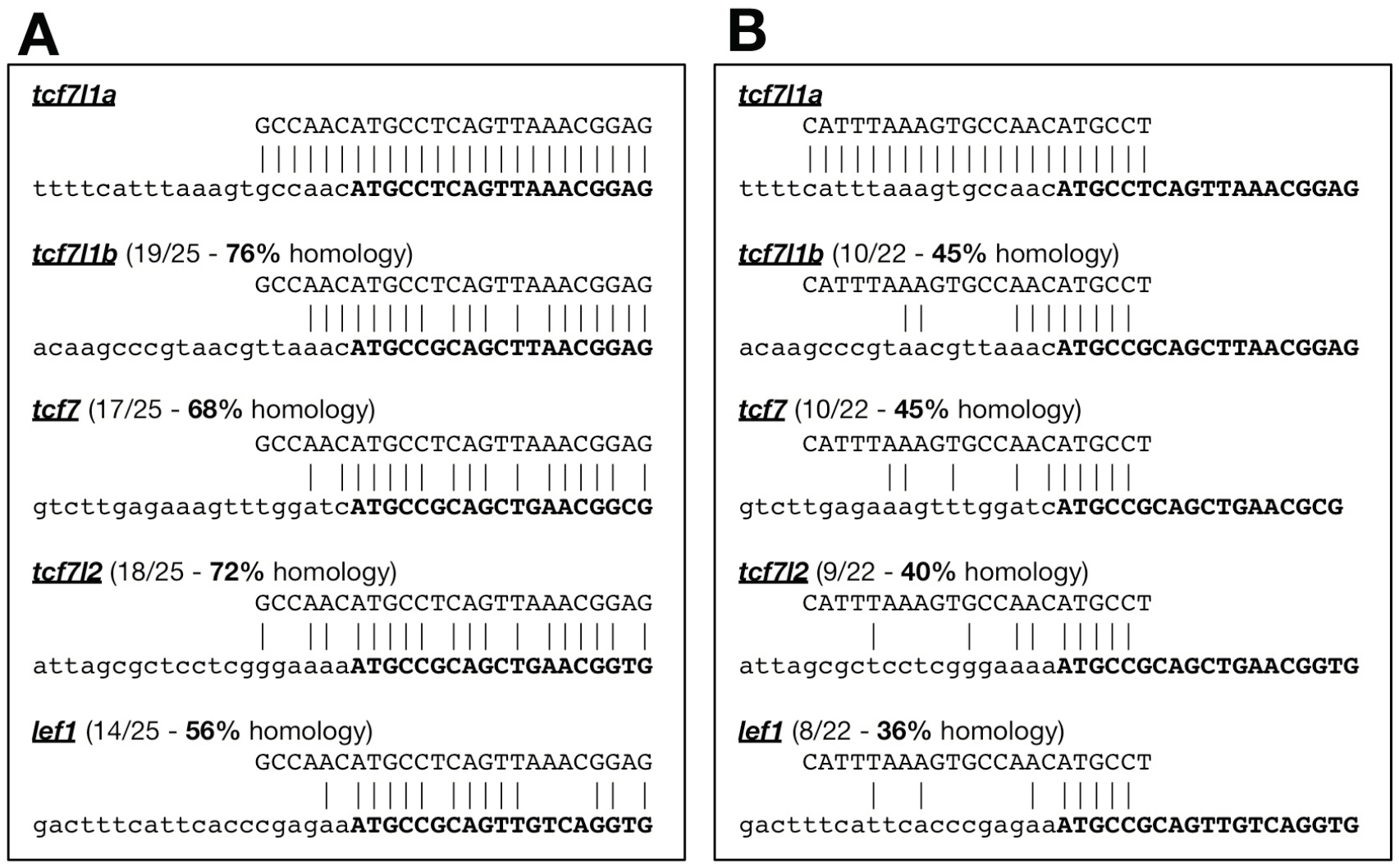
Alignment of *tcf7l1a,* morpholinos to *lef1* and other *tcf* genes. Alignment of the previously published *tcf7l1a* morpholino (**A**, mo1^*tcf7l1a*^, Dorsky et al. 2003), and mo2^*tcf7l1a*^ (**B**) with less homology to other *lef/tcf* genes. Target genes in bottom sequence, 5’UTR in lowercase, gene open reading frame in bold uppercase.

**Fig S3.**
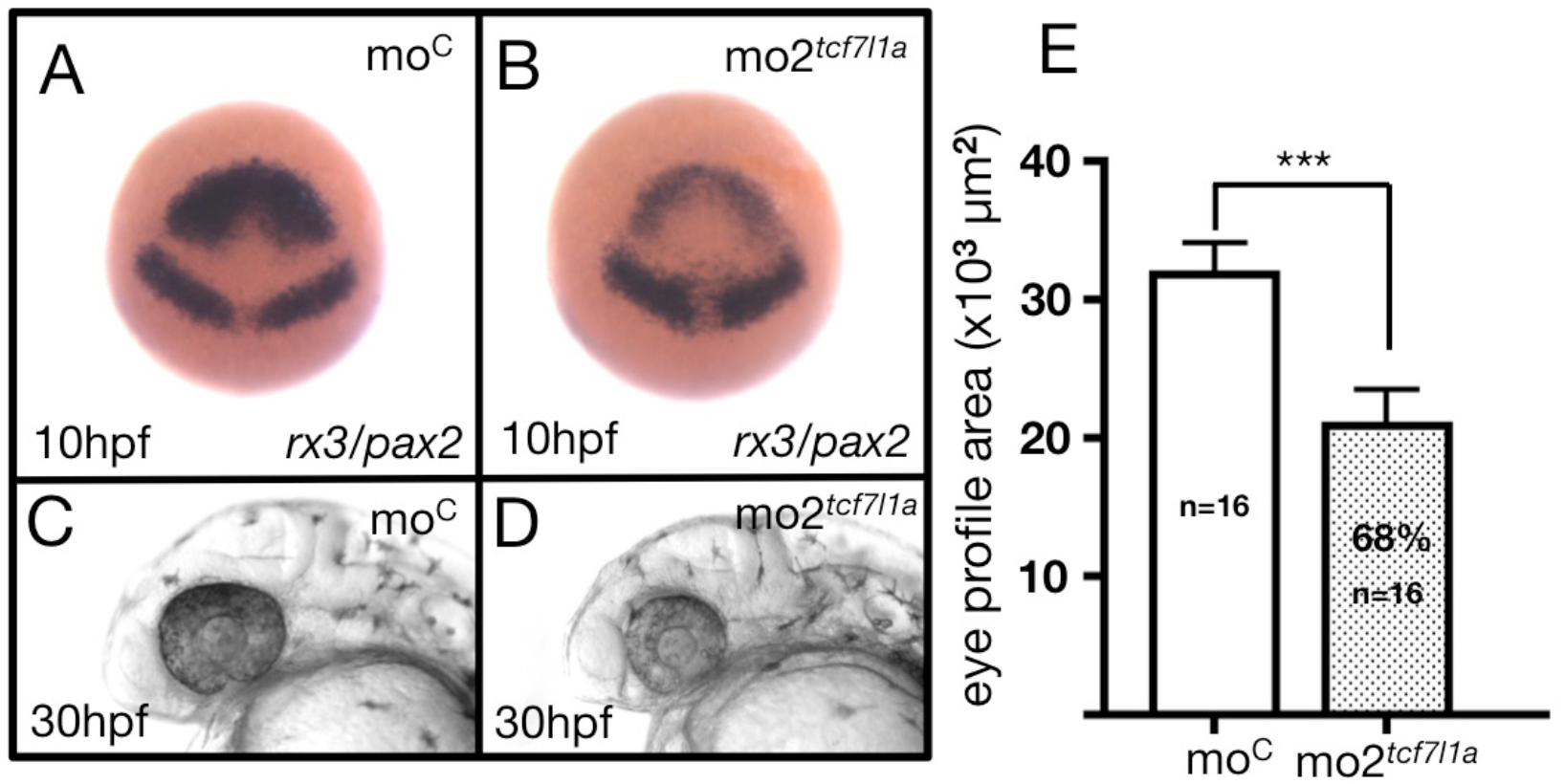
*tcf7l1a* morpholino (mo2^*tcf7l1a*^) phenocopies the *Ztcf7l1a*^*-/-*^ mutant. (**A, B**) Dorsal views of *rx3/pax2a in situ* hybridisation at 100%epibly and (**C, D**) lateral views of 30hpf wildtype embryos injected with 400pmols of (**A, C**) control morpholino or (**B, D**) _mo2_*tcf7l1a*. (**E**) Plot showing the quantification of the eye profile area in wildtype embryos injected with control morpholino (first bar) or mo2^*tcf7l1a*^ (second bar) at 30hpf.

**Fig S4.**
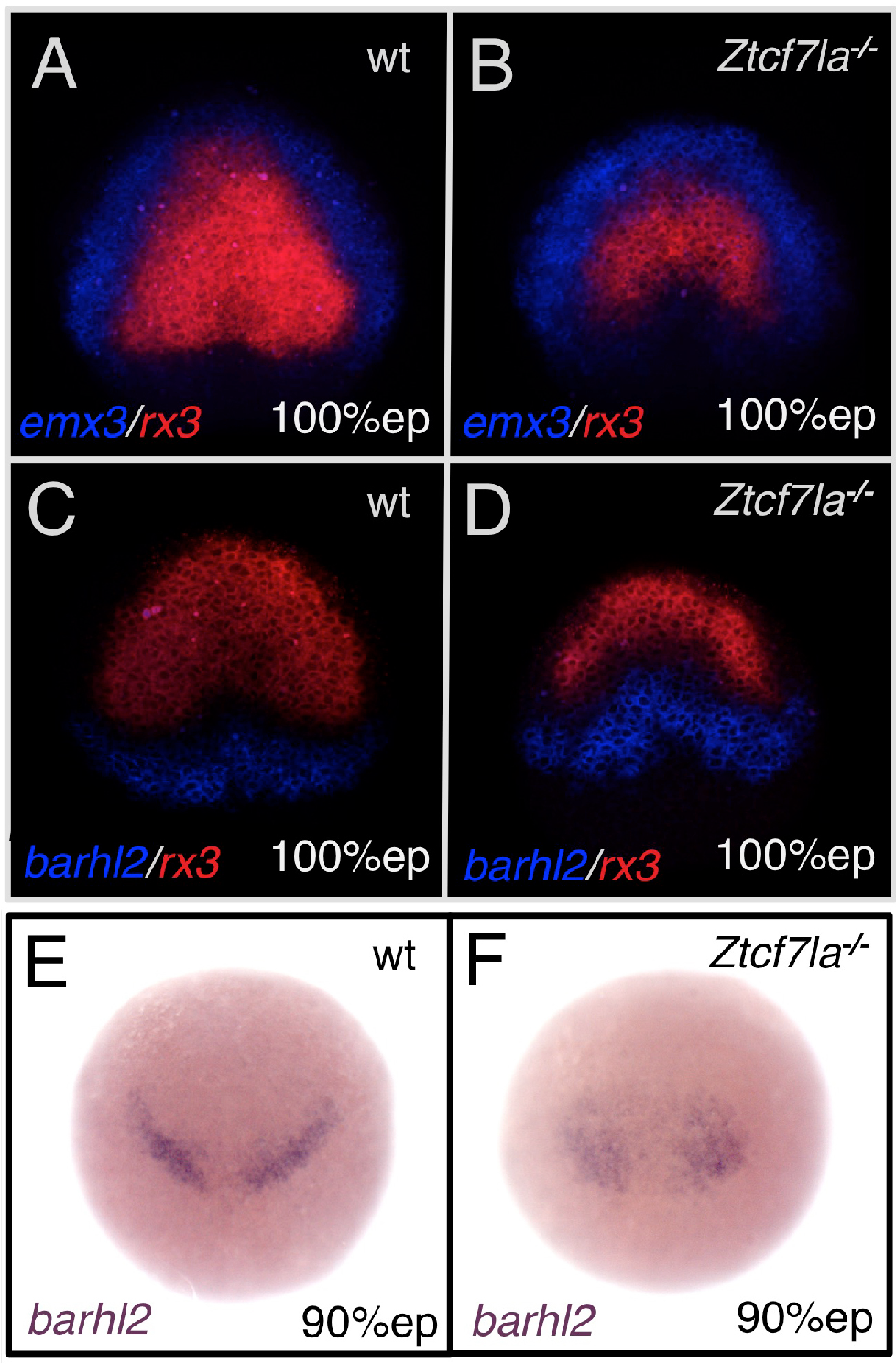
The eye field domain of the anterior neural plate is caudalised in *Ztcf7l1a* mutants. Dorsal views of anterior neural plates, anterior up. (**A-D**) Double fluorescent *in situ* hybridisation of prospective telencephalic marker *emx3* (blue) and prospective eye field marker *rx3* (red) (**A, B**), and prospective diencephalic marker *barhl2* (blue) and *rx3* (red) (**C, D**) in wildtype (**A, C**) and *Ztcf7l1a*^*-/-*^ (**B, D**) embryos at 10hpf. (**E, F**) *In situ* hybridisation of *barhl2* in wildtype (**E**) and *Ztcf7l1a*^*-/-*^ (**F**) embryos at 9hpf.

**Fig S5.**
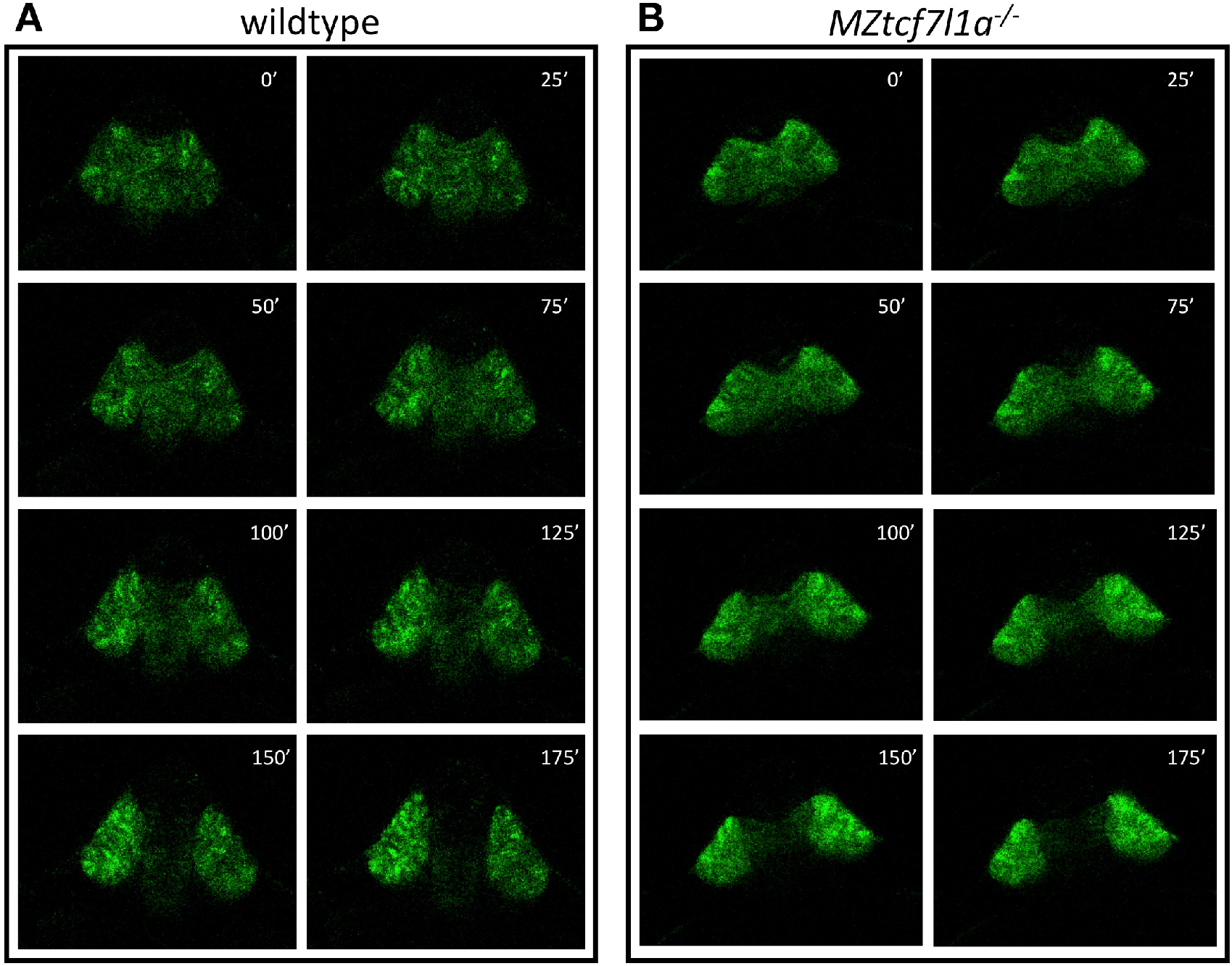
Eye vesicle evagination in heterozygous and *Ztcf7l1a*^*-/-*^ mutants. Confocal time lapse movie snapshots (1 frame every 25 minutes) of heterozygous sibling (**A**) and *Ztcf7l1a*^*-/-*^ mutants (**B**) expressing the *Tg(rx3:GFP)*^*zf460Tg*^ transgene. First frame taken at 11hpf. Minutes after movie has started indicated in each frame.

**Fig S6.**
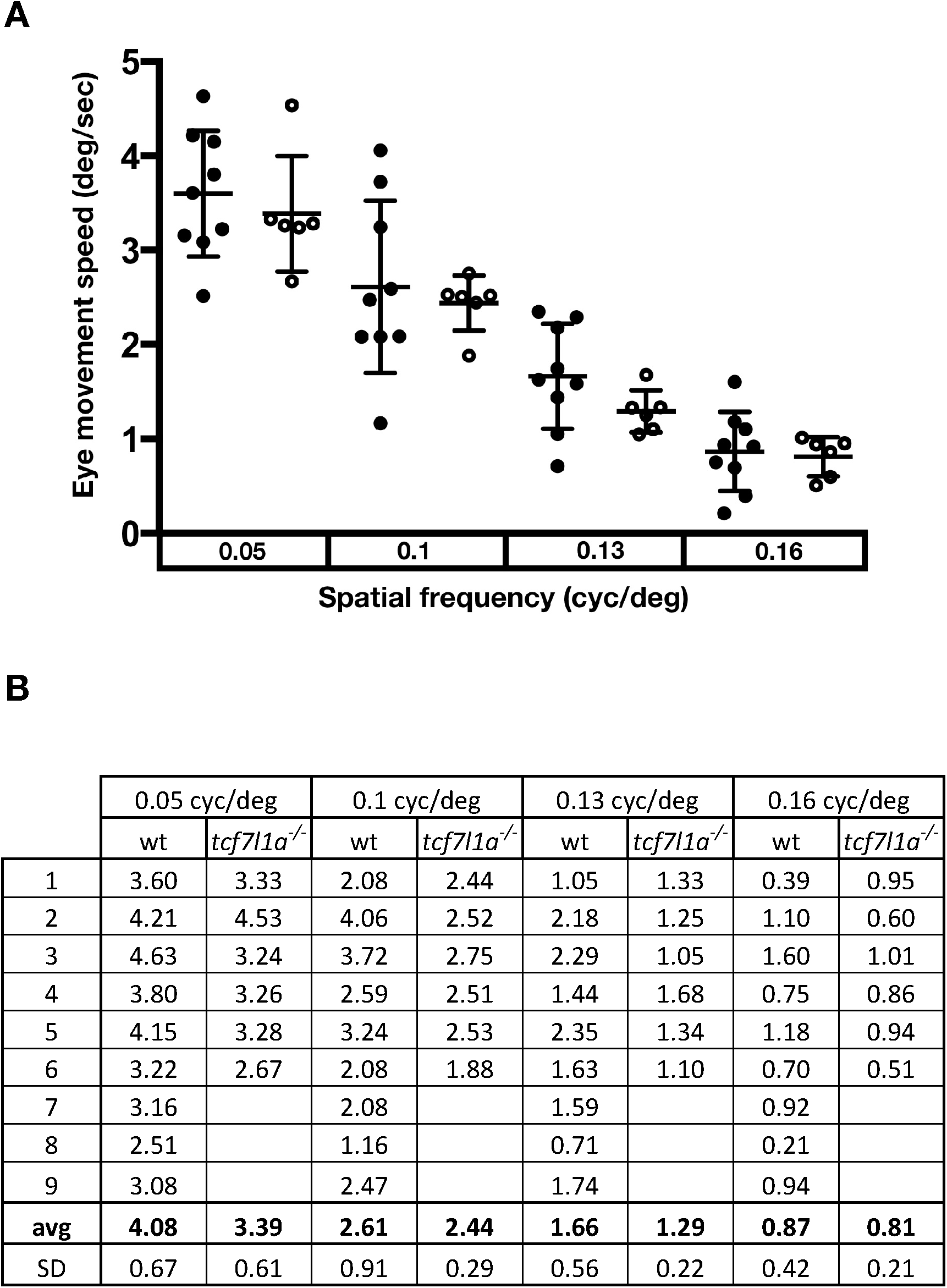
Optokinetic response analysis of wildtype and *Ztcf7l1a*^*-/-*^ mutants. (**A**) Average of left and right eye movement speed in degrees(deg)/second(sec), when wildtype (filled circles) and *Ztcf7l1a*^*-/-*^ mutants (empty circles) embryos were presented with vertical moving stripes at increasing angular speeds, 0.05, 0.1, 0.13 and 0.16 cycles(cyc)/deg. The data used to generate this plot is presented in TableS6. (**B**) Tabulation of the data used on the plot. No significant difference observed when performing an unpaired t-test with Welch’s correction. Average (avg), standard deviation (SD).

**Fig S7.**
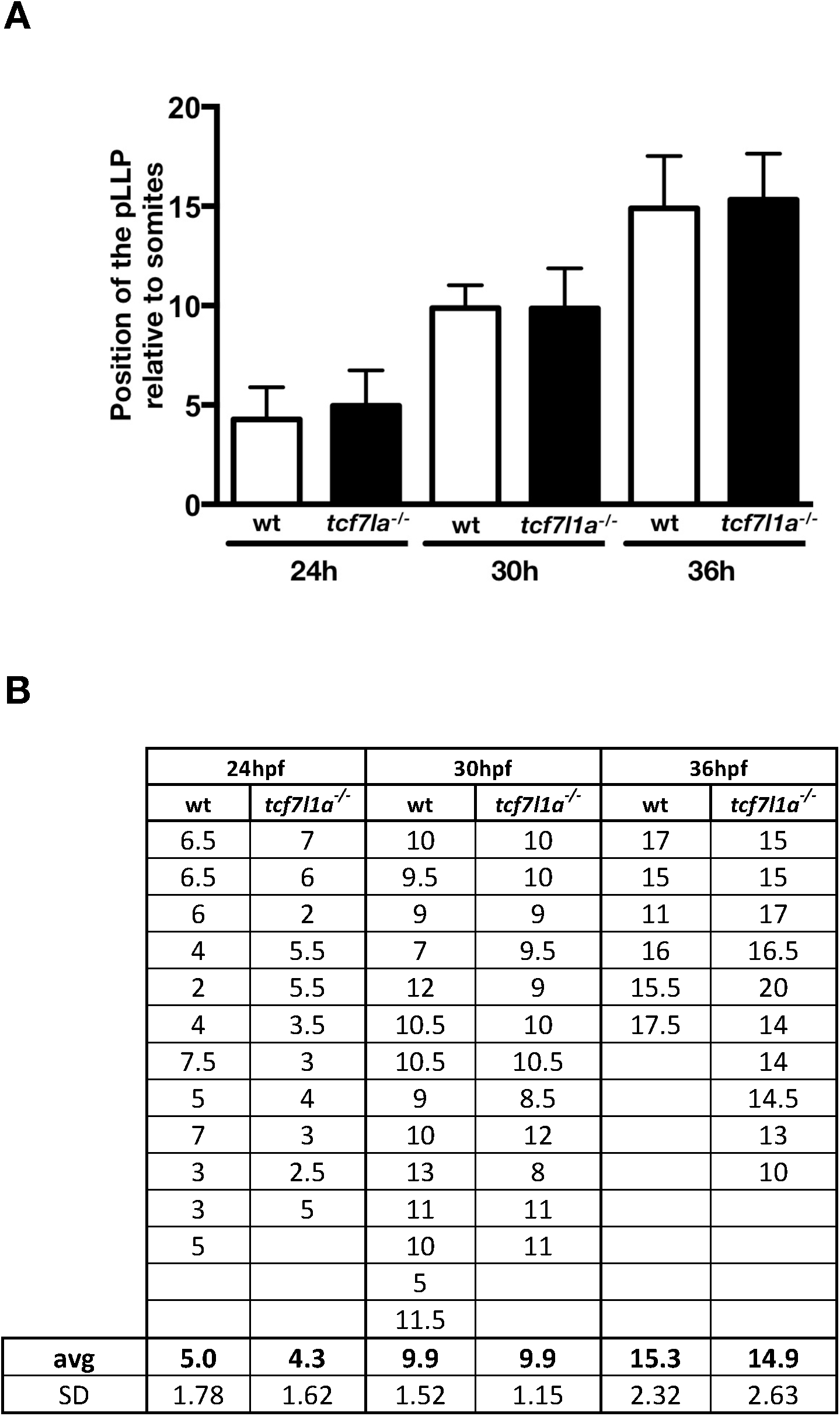
Quantification of the posterior lateral line primordium position in wildtype and *Ztcf7l1a*^*-/-*^ mutants. (**A**) Plot showing the position of the posterior lateral line primordium relative to somite number in wildtype (white bars) and *Ztcf7l1a*^*-/-*^ mutants (black bar) at 24, 30 and 36hpf. (**B**) Tabulation of the data used on the plot. No significant difference observed when performing an unpaired t-test with Welch’s correction. Average (avg), standard deviation (SD).

**Fig S8.**
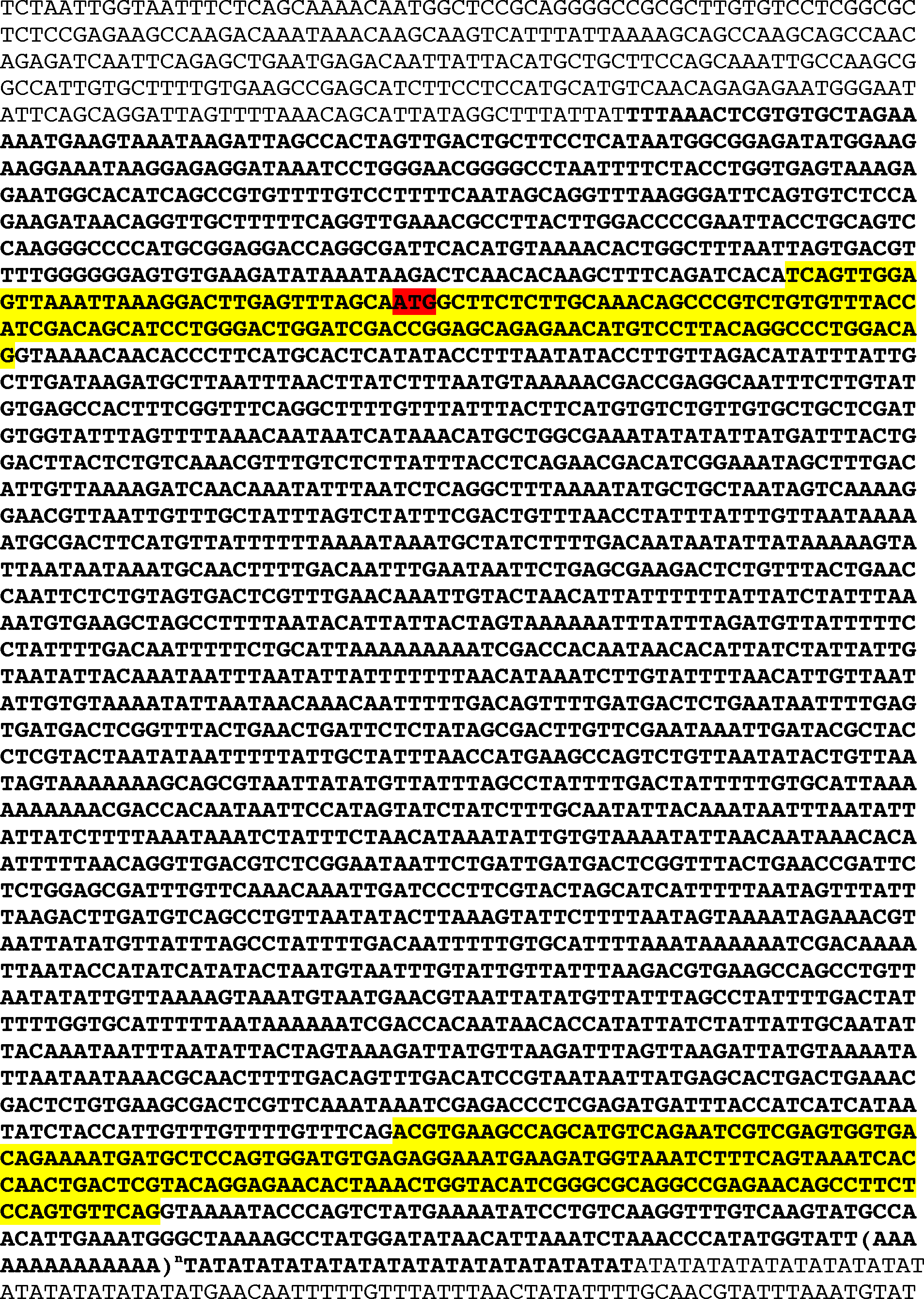
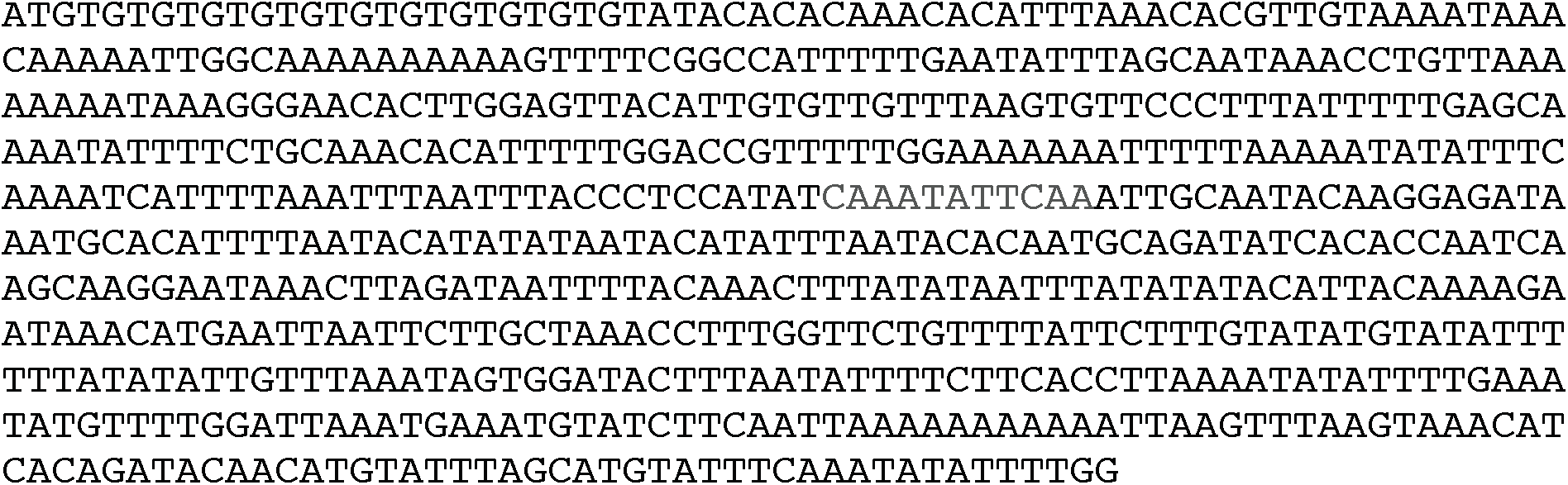
Genomic DNA sequence of *hesx1* locus spanning exons 1 and 2. Genomic DNA sequence deleted in U768 fish in bold. *hesx1* exons 1 and 2 highlighted in yellow. Open reading frame first codon in exon 1 is highlighted in red.

**Fig S9.**
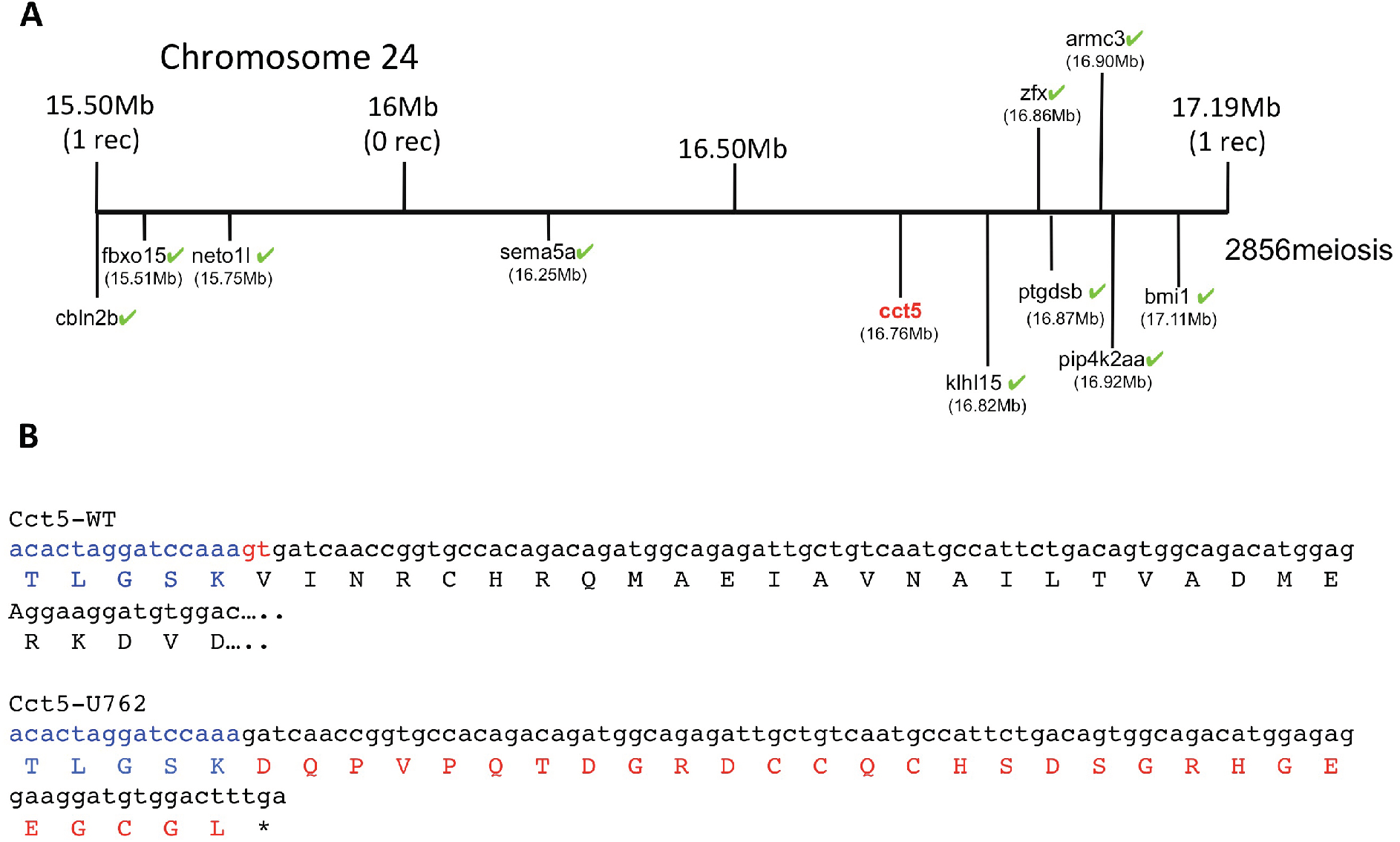
Genetic map of *U762* modifier of the *tcf7l1a* mutant phenotype and protein product of *cct5*^*U762*^. (**A**) Representation of the SSLP segregation linkage analysis mapping of ***U762*** modifier of *tcf71a* to a 1.69Mb interval on chromosome 12, between 15.50 Megabases (Mb) with 1 recombinant (rec) and 17.19Mb with 1 rec. Green ticks highlight sequenced genes in the interval that show no mutations. (**B**)Nucleotide and protein sequence of wildtype (top alignment) and *cct5*^*U762*^ (bottom alignment). *cct5 e*xon 4 nucleotides and amino acids in blue. The last two 3’ nucleotides in exon 4 that are used in *cct5*^*U762*^ as a splice donor in red. Nonsense amino acid sequence in *cct5*^*U762*^ in red.

**Table S1.** qRT-PCR data on mRNA levels from *Ztcf7l1a*^*-/-*^ versus wildtype embryos. Figures represent the fold change of *Ztcf7l1a*^*-/-*^ over wildtype qRT-PCR experiments performed as technical and biological triplicates using GAPDH as a reference control. Standard deviation (SD). P value calculated by an unpaired t-test with Welch’s correction.

|  | *lef1* | *tcf7* | *tcf7l1a* | *tcf7l1b* | *tcf7l2* | *otx1b* | *otx2* | *six3b* | *rx3* |
| --- | --- | --- | --- | --- | --- | --- | --- | --- | --- |
| sample1 | 0.95 | 0.9 | 0.18 | 0.65 | 0.7 | 0.89 | 0.93 | 0.35 | 0.24 |
| sample2 | 1.17 | 1.4 | 0.19 | 0.74 | 0.74 | 0.98 | 0.89 | 0.53 | 0.27 |
| sample3 | 0.67 | 0.75 | 0.12 | 0.5 | 0.43 | 0.55 | 0.57 | 0.45 | 0.28 |
| **average** | **0.93** | **1.02** | **0.16** | **0.63** | **0.62** | **0.81** | **0.80** | **0.44** | **0.26** |
| SD | 0.25 | 0.34 | 0.04 | 0.12 | 0.17 | 0.23 | 0.20 | 0.09 | 0.02 |
| p value | ns | ns | <0.0006 | <0.0341 | <0.058 | ns | ns | <0.091 | <0.0002 |

**Table S2.** Measurement of the volume of the prospective forebrain in wildtype and *Ztcf7l1a* mutants. Prospective forebrain size data generated by measuring the volume enclosed by *emx3* expression to the rostral limit of *pax2a* (mesencephalic marker) expression after whole mount in situ hybridisation in wildtype (+/+), *tcf7l1a*^*+/-*^ (+/-) and *Ztcf7l1a*^*-/-*^ (-/-) embryos at 10hpf. Data in μm^2^. Average (avg), standard deviation (SD). None significant, ns. P value calculated by an unpaired t-test with Welch’s correction.

| | *emx3/pax2* enclosed area ($\mu\text{m}^2$) | | |
| --- | --- | --- | --- |
| *tcf7l1a* | +/+ | +/- | -/- |
| 1 | 57356 | 58827 | 41345 |
| 2 | 68208 | 50347 | 41664 |
| 3 | 46505 | 77337 | 45372 |
| 4 | 57387 | 50698 | 40291 |
| 5 | 55112 | 59242 | 44298 |
| 6 | 58241 | 59966 | 42625 |
| 7 | 65606 | 59620 | 39436 |
| 8 | 53148 | 57212 | 40810 |
| 9 |  | 57569 | 47467 |
| 10 |  | 68588 | 40511 |
| 11 |  | 53195 | 61870 |
| 12 |  | 66222 |  |
| 13 |  | 50361 |  |
| 14 |  | 60196 |  |
| 15 |  | 51393 |  |
| **avg** | **57695** | **58718** | **44154** |
| SD | 6826 | 7546 | 6359 |
| p value |  | ns | >0.0004 |

**Table S3.**
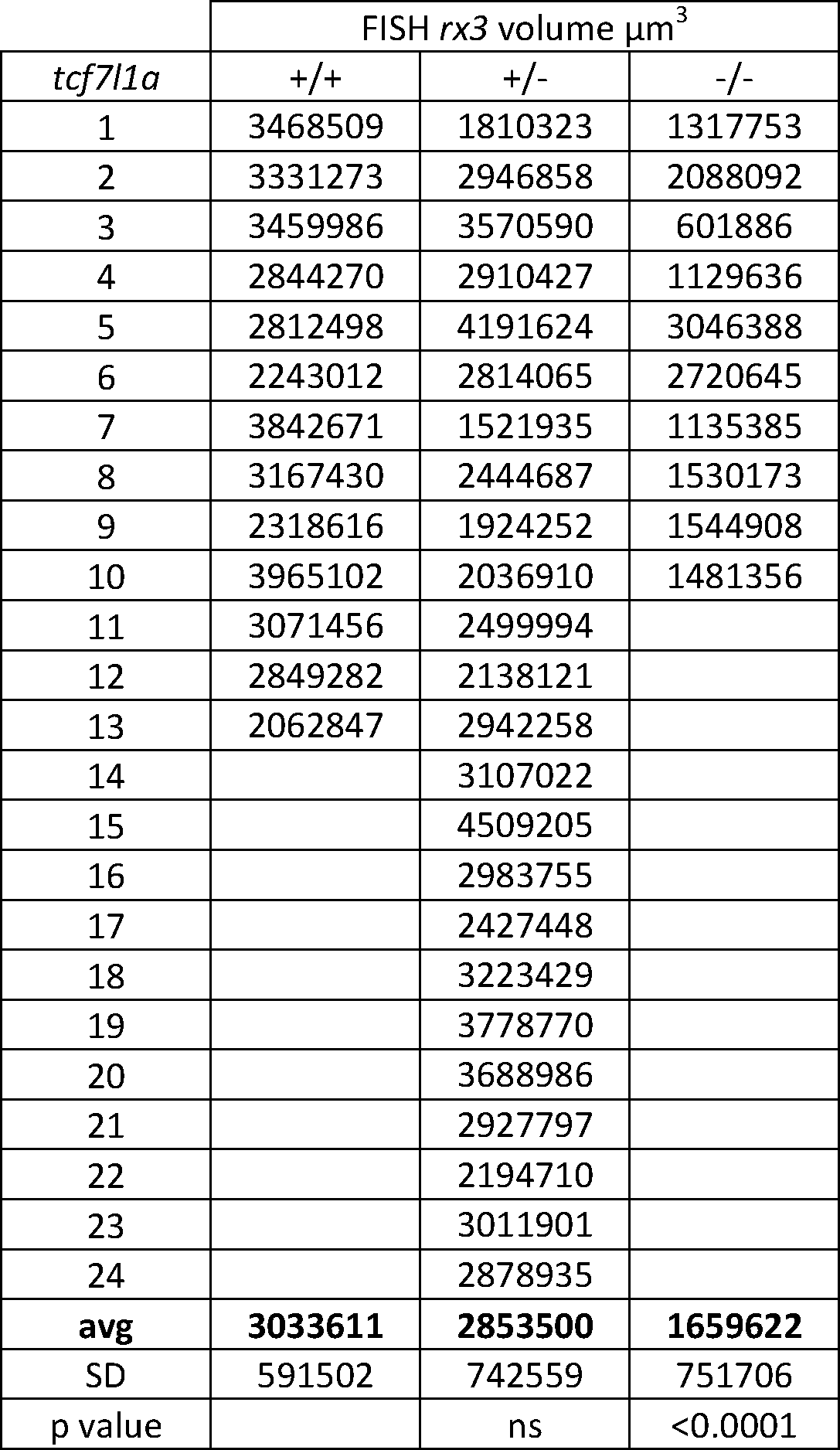
Measurement of the volume of *rx3* expression in the eye field by fluorescent *in situ* hybridisation in wildtype and *tcf7l1a* mutants. Confocal volume analysis of *rx3* fluorescent in situ hybridisation (FISH) on wildtype (+/+), *tcf7l1a*^*+/-*^ (+/-) and *Ztcf7l1a*^*-/-*^ (-/-) at 10hpf. Data in μm^3^. Average (avg), standard deviation (SD). none significant (ns). P value calculated by an unpaired t-test with Welch’s correction.

| | FISH *rx3* volume $\mu\text{m}^3$ | | |
| --- | --- | --- | --- |
| *tcf7l1a* | +/+ | +/- | -/- |
| 1 | 3468509 | 1810323 | 1317753 |
| 2 | 3331273 | 2946858 | 2088092 |
| 3 | 3459986 | 3570590 | 601886 |
| 4 | 2844270 | 2910427 | 1129636 |
| 5 | 2812498 | 4191624 | 3046388 |
| 6 | 2243012 | 2814065 | 2720645 |
| 7 | 3842671 | 1521935 | 1135385 |
| 8 | 3167430 | 2444687 | 1530173 |
| 9 | 2318616 | 1924252 | 1544908 |
| 10 | 3965102 | 2036910 | 1481356 |
| 11 | 3071456 | 2499994 |  |
| 12 | 2849282 | 2138121 |  |
| 13 | 2062847 | 2942258 |  |
| 14 |  | 3107022 |  |
| 15 |  | 4509205 |  |
| 16 |  | 2983755 |  |
| 17 |  | 2427448 |  |
| 18 |  | 3223429 |  |
| 19 |  | 3778770 |  |
| 20 |  | 3688986 |  |
| 21 |  | 2927797 |  |
| 22 |  | 2194710 |  |
| 23 |  | 3011901 |  |
| 24 |  | 2878935 |  |
| **avg** | **3033611** | **2853500** | **1659622** |
| SD | 591502 | 742559 | 751706 |
| p value |  | ns | <0.0001 |

**Table S4.** Eye volume measurement data in wildtype and *tcf7l1a* mutants. Data of eye volume measurement figures in μm^3^ in wildtype and *tcf7l1a*^*-/-*^ embryos at 24, 28, 32, 36, 48, 60, 72 and 96hpf. Average (avg), standard deviation (SD), percentage of *Ztcf7l1a*^*-/-*^ eye volume size relative to wildtype eyes (%).

|  | 24hpf |  | 28hpf |  | 32hpf |  | 36hpf |  |
| --- | --- | --- | --- | --- | --- | --- | --- | --- |
|  | wt | *tcf7l1a*<sup>-/-</sup> | wt | *tcf7l1a*<sup>-/-</sup> | wt | *tcf7l1a*<sup>-/-</sup> | wt | *tcf7l1a*<sup>-/-</sup> |
|  | 2173725 | 1017791 | 2342908 | 1576503 | 2451428 | 1828463 | 3658209 | 2460898 |
|  | 2303076 | 1051526 | 2570537 | 1917720 | 2489175 | 2039707 | 3274603 | 2439886 |
|  | 1705511 | 1066121 | 2337095 | 1070995 | 2845175 | 1978270 | 3207975 | 2232993 |
|  | 1955287 | 1308467 | 2348931 | 1790742 | 3006924 | 1914276 | 2897337 | 2221198 |
|  | 2805372 | 1326337 | 2351628 | 1725672 | 2842500 | 1930381 | 3303325 | 2015056 |
|  | 2253508 | 1301159 | 2735274 | 1637355 | 2785130 | 1741815 | 2697041 | 3066122 |
|  | 1774861 | 1328035 | 2467551 | 1604776 | 2551566 | 1841438 | 3137085 | 1800979 |
|  | 2487443 | 1301917 | 2672275 | 1617131 | 2999859 | 1947938 | 3384282 | 2323963 |
|  | 1718721 | 1403183 | 2673491 | 1404137 | 2851039 | 2218943 | 3367798 | 2622681 |
|  | 2255991 | 1641569 | 2510968 | 1856147 | 3378945 | 1862151 | 3615953 | 2352265 |
|  | 2043767 | 1135015 | 2495736 | 1966110 | 2917531 | 2044091 | 3122494 | 2814747 |
|  | 2630277 | 917394 | 2476897 | 1459703 | 3067765 | 2049230 | 2921844 | 2511973 |
|  |  | 1094766 | 2572070 |  | 3020867 | 2289502 | 3046882 | 2860238 |
|  |  | 1538292 | 2287218 |  |  | 1665499 | 3359380 | 2701119 |
|  |  |  | 2632288 |  |  | 1894067 | 2860411 |  |
|  |  |  |  |  |  |  | 3447190 |  |
| avg | 2175628 | 1245112 | 2498324 | 1635583 | 2862146 | 1949718 | 3206363 | 2458866 |
| SD | 354914 | 206675 | 142882 | 248218 | 256341 | 164423 | 272999 | 341047 |
| % | 57.2 |  | 65.5 |  | 68.1 |  | 76.7 |  |

|  | 48hpf |  | 60hpf |  | 72hpf |  | 96hpf |  |
| --- | --- | --- | --- | --- | --- | --- | --- | --- |
|  | wt | *tcf7l1a*<sup>-/-</sup> | wt | *tcf7l1a*<sup>-/-</sup> | wt | *tcf7l1a*<sup>-/-</sup> | wt | *tcf7l1a*<sup>-/-</sup> |
|  | 3999209 | 3307657 | 4100360 | 3883270 | 5009654 | 4145880 | 5455460 | 5167760 |
|  | 4497099 | 3004106 | 4984899 | 4002063 | 4880009 | 4385725 | 6215474 | 5465572 |
|  | 4266652 | 3720744 | 4516578 | 4147457 | 4793584 | 3970719 | 6661246 | 4610702 |
|  | 4353610 | 2987478 | 4531155 | 3651250 | 5259776 | 3960662 | 6229111 | 5724542 |
|  | 4411489 | 4023028 | 3879568 | 3763464 | 4788932 | 4331574 | 5666545 | 5198084 |
|  | 4251961 | 4151421 | 4491768 | 3736227 | 5238466 | 4602247 | 6315093 | 5364821 |
|  | 4041664 | 3767914 | 4631971 | 4491069 | 4934159 | 4039358 | 5973419 | 5764363 |
|  | 4278292 | 3633181 | 4396588 | 3982956 | 5007355 | 4400128 | 6358613 | 4601883 |
|  | 3280615 | 3766971 | 4608893 | 3390604 | 5117971 | 4641026 | 6503686 | 5465686 |
|  | 4567178 | 3082474 | 4837262 | 3372391 | 5046112 |  | 6611962 | 5134945 |
|  | 4309538 | 3648767 | 4631242 | 3638188 | 5140014 |  | 6058679 | 5523381 |
|  |  | 3437048 |  | 3811614 | 5155990 |  | 5635639 | 4815731 |
|  |  | 4451579 |  | 4351661 | 5057506 |  | 6161109 | 5278184 |
|  |  | 3826961 |  | 4136280 |  |  |  | 5251031 |
|  |  | 3641260 |  |  |  |  |  | 5092468 |
|  |  | 3136116 |  |  |  |  |  |  |
|  |  | 3467977 |  |  |  |  |  |  |
|  |  | 3627969 |  |  |  |  |  |  |
|  |  | 3923272 |  |  |  |  |  |  |
| avg | 4205210 | 3610838 | 4510026 | 3882750 | 5033041 | 4275258 | 6142003 | 5230610 |
| SD | 350291 | 393089 | 308370 | 328796 | 152611 | 258790 | 376463 | 351597 |
| % | 85.9 |  | 86.1 |  | 84.9 |  | 85.2 |  |

**Table S5.** Classification and quantification of *atoh7* expression patterns in wildtype and *Ztcf7l1a*^*-/-*^ eyes. Quantification of *atoh7* expression categories in the eye at 28, 32, 36, 40, 44, 48 and 52hpf in wildtype (top table) and *Ztcf7l1a*^*-/-*^ (bottom table) embryos. *atoh7* expression was classified in the following categories: VN, ventro nasal; VN+, ventro nasal plus a few scattered cells; N+, nasal plus scattered cells covering the whole retina; NR, nasal retina; WR, whole retina; PR, peripheral retina.

| *tcf7l1a*<sup>+/+</sup> | VN | VN+ | N+ | WR | PR | total |
| --- | --- | --- | --- | --- | --- | --- |
| 28hpf | 0 | 17(61%) | 11(39%) | 0 | 0 | 28 |
| 32hpf | 0 | 0 | 7(26%) | 20(74%) | 0 | 27 |
| 36hpf | 0 | 0 | 0 | 12(34%) | 23(66%) | 35 |
| 40hpf | 0 | 0 | 0 | 4(13%) | 26(87%) | 30 |
| 44hpf | 0 | 0 | 0 | 0 | 25(100%) | 25 |
| 48hpf | 0 | 0 | 0 | 0 | 30(100%) | 30 |
| 52hpf | 0 | 0 | 0 | 0 | 32(100%) | 32 |

| *tcf7l1a*<sup>-/-</sup> | VN | VN+ | NR | WR | PR | total |
| --- | --- | --- | --- | --- | --- | --- |
| 28hpf | 10(91%) | 1(9%) | 0 | 0 | 0 | 11 |
| 32hpf | 5(42%) | 7(58%) | 0 | 0 | 0 | 12 |
| 36hpf | 0 | 4(44%) | 5(56%) | 0 | 0 | 9 |
| 40hpf | 0 | 0 | 4(36%) | 7(64%) | 0 | 11 |
| 44hpf | 0 | 0 | 0 | 12(100%) | 0 | 12 |
| 48hpf | 0 | 0 | 0 | 7(64%) | 4(36%) | 11 |
| 52hpf | 0 | 0 | 0 | 0 | 8(100%) | 8 |

**Table S6.**
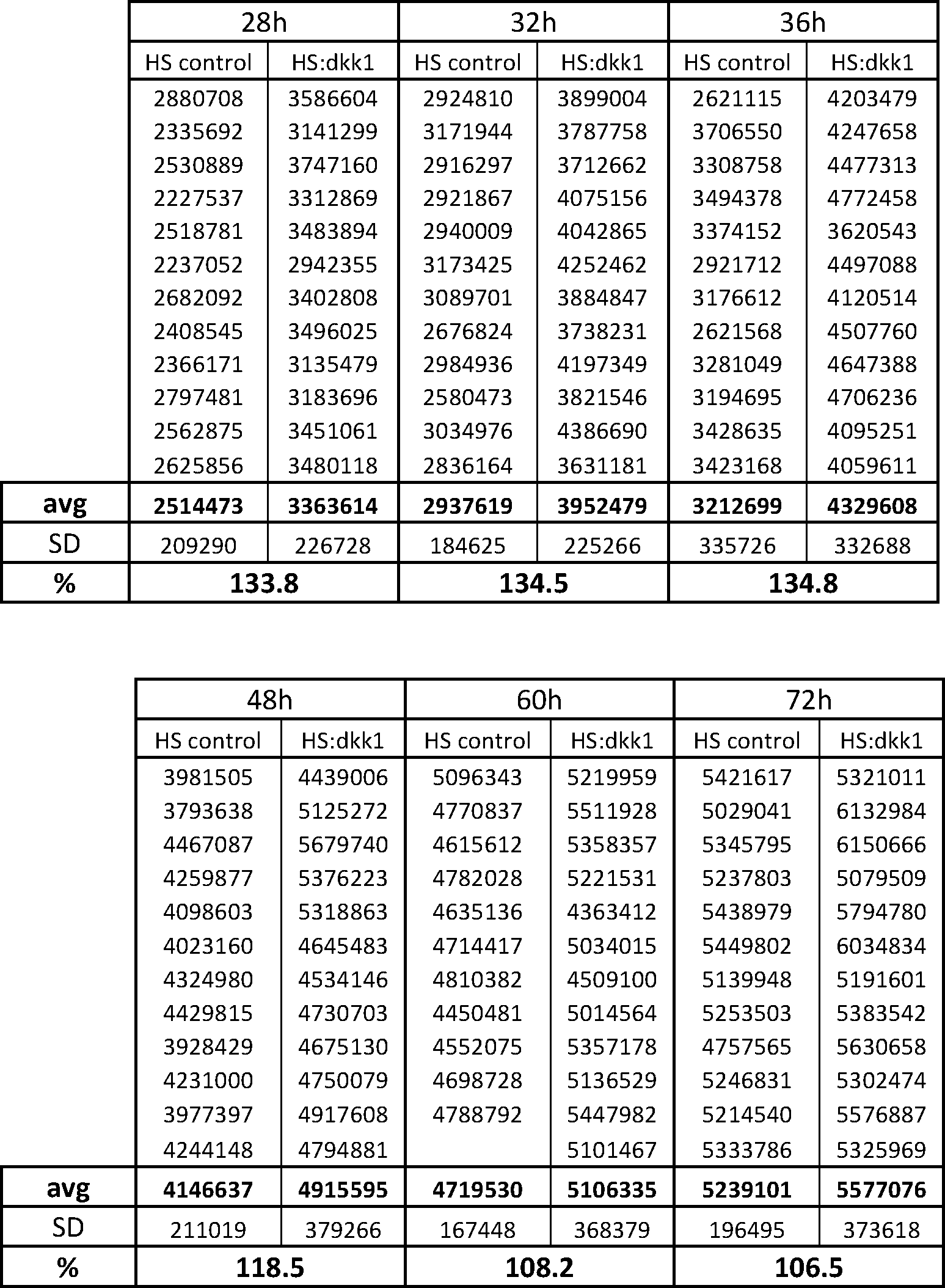
Eye volume measurement data in heat-shocked control wildtype and *Tg(HS:dkk1)*^*w32*^ embryos. Tabulation of eye volume measurements in μm^3^ in heat shock control wild-type and *Tg(HS:dkk1)*^*w32*^ embryos at 28, 32, 36, 48, 60 and 72, used for plot in Fig.5R. Average (avg), standard deviation (SD), percentage of *Tg(HS:dkk1)*^*w32*^ eye volume size relative to heat shock control wildtype eyes (%).

**Table S7.**
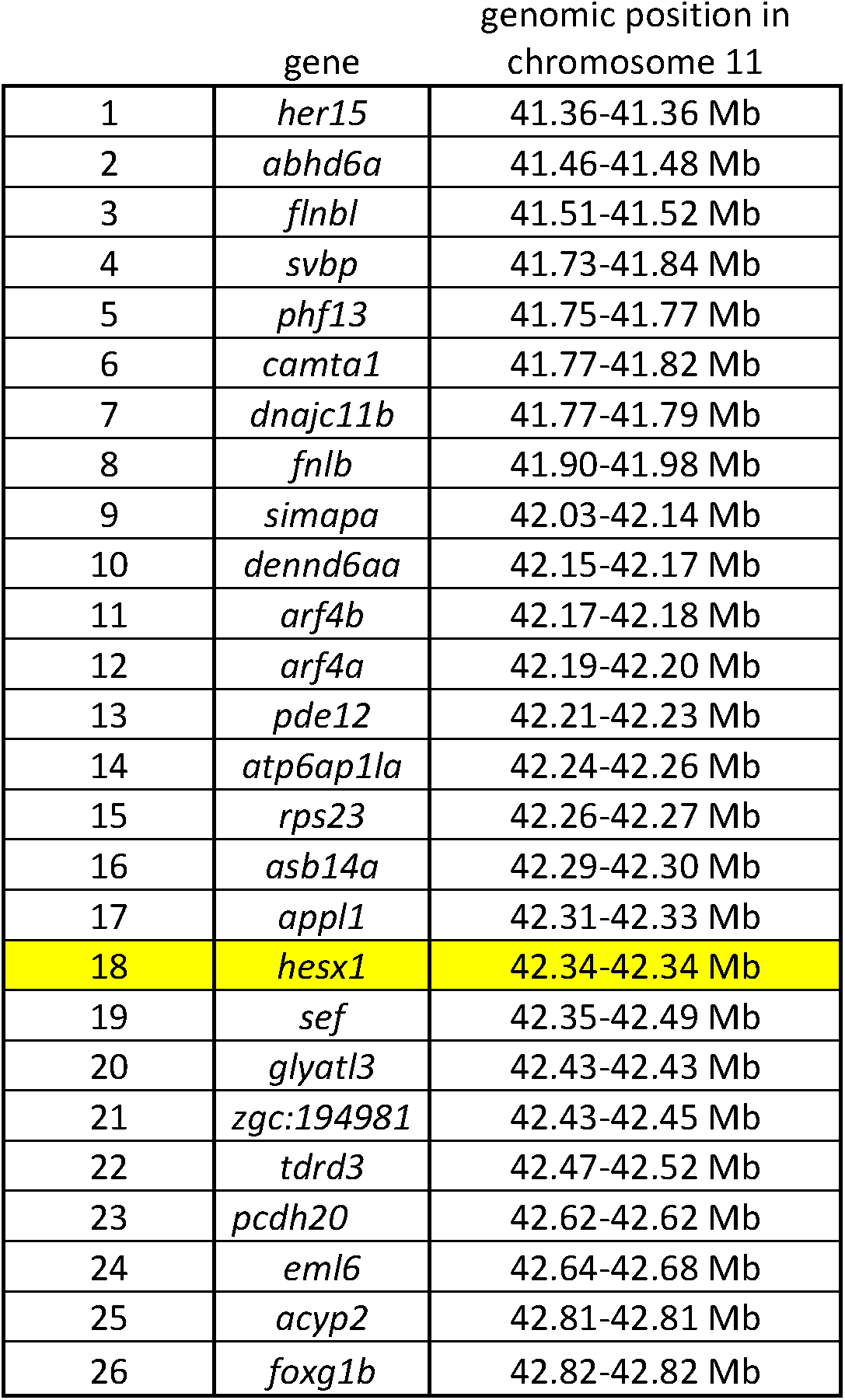
List of genes in the *U768* genetic interval. Position of the genes in the *U768* interval in megabases (Mb) in the GRCz10 assembly. Mapped gene is highlighted in yellow.

**Table S8.**
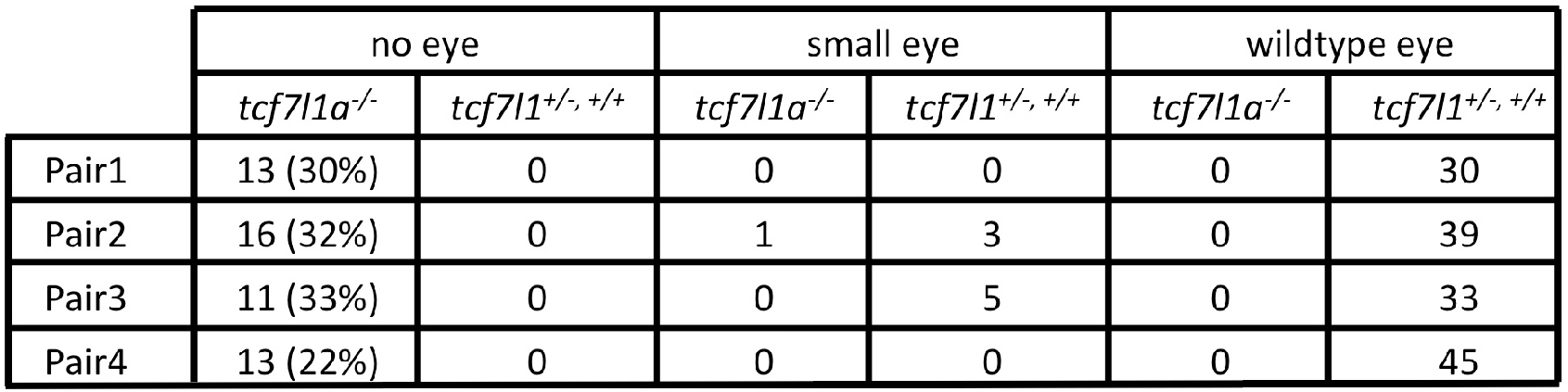
Frequency of eyeless embryos and their respective genotypes in four incrosses of *hesx1*^*-/-*^*/tcf7l1a*^*+/-*^ fish.

**Table S9.**
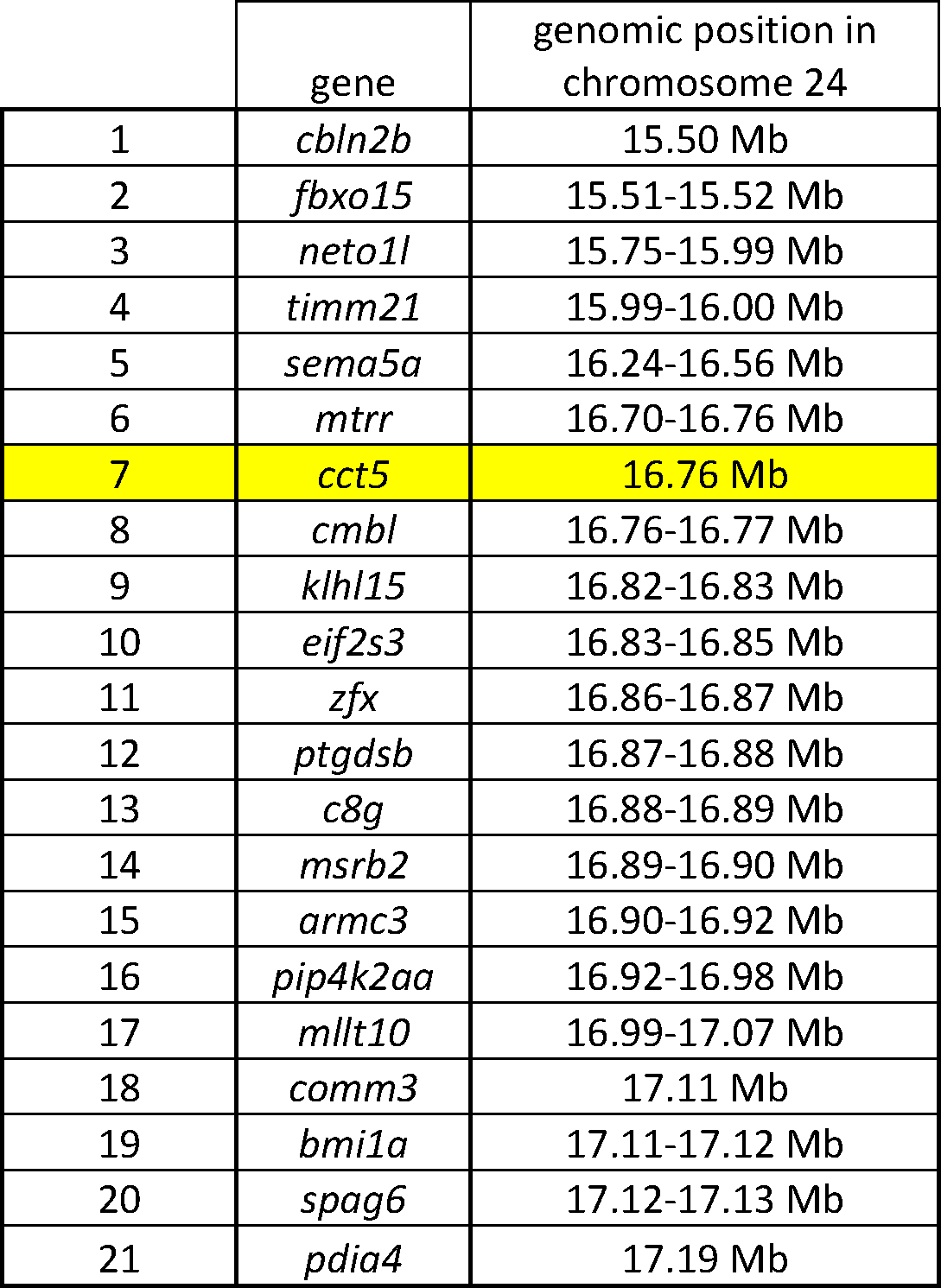
List of the genes in the *U762* genetic interval. Position of genes in the *U762* interval in megabases (Mb) in the GRCz10 assembly. Mapped gene is highlighted in yellow.

**Table S10.** Eye volume measurement data in wildtype, *tcf7l1a, gdf6a* and *gdf6a/tcf7l1a* double mutant sibblings. Data of eye volume measurement figures in μm^3^ in embryos at 80hpf. Average (avg), standard deviation (SD), percentage of mutant eye volume relative to wildtype eyes (%).

|  | wt | *tcf7l1a*<sup>-/-</sup> | *gdf6a*<sup>-/-</sup> | *gdf6a/tcf7l1a*<sup>-/-</sup> |
| --- | --- | --- | --- | --- |
|  | 2903277 | 2787301 | 2008253 | 1264302 |
|  | 2819887 | 2299871 | 2128333 | 1119602 |
|  | 3008809 | 2718549 | 2274431 | 1159715 |
|  | 2961193 | 2807418 | 2505976 | 1564565 |
|  | 2890943 |  | 2547329 | 1496518 |
|  | 3147872 |  |  | 1364723 |
|  | 2701579 |  |  |  |
|  | 3100625 |  |  |  |
| avg | 2941773 | 2653285 | 2292864 | 1328238 |
| SD | 145993 | 238661 | 233764 | 179731 |
| % |  | 90 | 78 | 45 |

**Movies S1 and S2. Time lapse movies of eye vesicle evagination in heterozygous sibling and *Ztcf7l1a*^*-/-*^ mutants.**

Confocal time lapse movies (1 frame every 5 minutes) of heterozygous sibling (**S1**) and *Ztcf7l1a*^*-/-*^ mutants (**S2**) expressing the *Tg(rx3:GFP)*^*zf460Tg*^ transgene. First frame taken at 11hpf; membrane RFP in red counterstain.

